# Monomeric α-Synuclein activates the Plasma Membrane Calcium Pump

**DOI:** 10.1101/2022.02.21.481193

**Authors:** Antoni Kowalski, Cristine Betzer, Sigrid Thirup Larsen, Emil Gregersen, Estella Anne Newcombe, Montaña Caballero Bermejo, Annette Eva Langkilde, Birthe Brandt Kragelund, Poul Henning Jensen, Poul Nissen

**Affiliations:** Department of Molecular Biology and Genetics, Aarhus University, Aarhus, Denmark; Department of Biomedicine, Aarhus University, Aarhus, Denmark; Danish Research Institute of Translational Neuroscience – DANDRITE, Aarhus University, Aarhus, Denmark; Department of Drug Design and Pharmacology, University of Copenhagen, Denmark; REPIN and Structural Biology and NMR Laboratory, Department of Biology, University of Copenhagen, Denmark; Department Biochemistry and Molecular Biology and Genetics, IBMP, University of Extremadura, Badajoz, Spain

## Abstract

Alpha-synuclein (aSN) is a membrane-associated and intrinsically disordered protein, well-known for pathological aggregation in neurodegeneration. The physiological function of aSN however is disputed. Pull-down experiments have pointed to plasma membrane Ca^2+^-ATPase (PMCA) as a potential interaction partner. From proximity ligation assays we find that aSN and PMCA colocalize at neuronal synapses, and that calcium expulsion is activated by aSN and PMCA. From PMCA activity studies we show that soluble, monomeric aSN activates PMCA at par with CaM, yet independent of the autoinhibitory domain of PMCA, but highly dependent on acidic phospholipids and membrane-anchoring of aSN. On PMCA, the key site is mapped to the acidic lipid-binding site, located within a disordered PMCA-specific loop region connecting the cytosolic A domain and transmembrane segment 3. Our studies point towards a novel physiological role of monomeric aSN as a stimulator of calcium clearance in neurons through activation of PMCA.

## INTRODUCTION

Alpha-synuclein (aSN, 140 residues, and 14.5 kDa molecular weight) is an intrinsically disordered protein, well recognized and highly studied for its pathological role in neurodegeneration. Oligomerization, fibrillization, and abnormal aggregation in neurons are linked to synucleinopathies, e.g., Lewy body dementia or Parkinson’s disease (PD) (Burre *et al*, 2018; Oliveira *et al*, 2021). However, the normal function of monomeric aSN remains an open question. Importantly, aSN is highly abundant in the presynaptic region of neurons, where concentrations reach 5-50 *μ*M (Bodner *et al*, 2009; Perni *et al*, 2017; Theillet *et al*, 2016). Even at this level, the protein remains structurally disordered and monomeric in cells (Fauvet *et al*, 2012; Theillet *et al*., 2016). Alpha-synuclein is also a peripheral membrane protein adopting a helical structure in the positively charged N-terminal region 1-95 through interaction with acidic phospholipids (Dikiy & Eliezer, 2012). The ability of aSN to interact with membranes is important for functions such as clustering of synaptic vesicles (Jo *et al*, 2000; Lautenschlager *et al*, 2018), regulation of the presynapse size (Vargas *et al*, 2017), and neurotransmitter release by promoting the formation of SNARE complexes (Burre *et al*, 2010). A disease-related mutation A30P – strongly linked to the inherited form of PD – results in diminished lipid-binding properties (Jensen *et al*, 1998). The C-terminal region, comprising residues 96-140 remains disordered regardless of interactions, has a strong negative charge, and binds calcium ions with low affinity (Eliezer *et al*, 2001; Lautenschlager *et al*., 2018; Nielsen *et al*, 2001).

Regulation of intracellular calcium homeostasis is essential for the proper functioning of cells. Resting concentration in the cytosol of a healthy cell is low, around 100 nM, while the extracellular concentration is in the millimolar range. In a calcium signaling event, rapid calcium influxes via calcium channels must be followed by efficient recovery by specialized proteins. The key players are i) the plasma membrane calcium ATPase (PMCA) and ii) the sodium-calcium exchanger (NCX), both removing Ca^2+^ to the extracellular space, and iii) the sarco/endoplasmatic reticulum Ca2+-ATPase (SERCA) filling intracellular calcium stores. Furthermore, calcium-binding proteins, in particular calmodulin (CaM), function as a calcium buffer and calcium-dependent regulator of multiple proteins (Cali *et al*, 2018).

Increasing evidence reveals a link between calcium dysregulation and propagation of synucleinopathies. Voltage-gated Ca_V_1 channels and Ca_V_1.3 mRNA are upregulated as an early feature of PD in areas not associated with overt loss of neurons or Lewy body formation (Hurley *et al*, 2013; Hurley *et al*, 2015). Fibrillar oligomers of aSN, which are formed with the disease development, can enhance the Ca^2+^-permeability of plasma membrane (Cali *et al*, 2014; Di Scala *et al*, 2016; Rcom-H’cheo-Gauthier *et al*, 2016) and have been found to activate SERCA (Betzer *et al*, 2018), thus contributing to disturbances in calcium homeostasis. The SERCA interaction was identified by aSN pull-down experiments, and PMCA was also identified from these experiments (Betzer *et al*, 2015).

Neurons have a large complexity and a highly polarized architecture; calcium signaling events in these cells, therefore, are highly localized. Dramatic, but extremely local calcium influxes are related to signals leading to vesicular exocytosis in the presynaptic termini. In the presynaptic bouton, the signaling events take place in nanodomains within 50 nm from Ca^2+^ channels (Augustine *et al*, 2003) and free calcium concentrations can rise locally by >1000-fold (Long *et al*, 2008). Terminating, rather than dissipating a local signal causes a high demand for an efficient and flexible calcium removal system with fine-tuning to a required level of resting concentrations.

PMCA is a single polypeptide transmembrane protein of 130-140 kDa and belongs to the P2B subfamily of P-type ATPases. Being a high affinity and low-capacity active transporter, it can fine-tune the resting free calcium ion concentration in cytosol (Carafoli, 1994; Strehler *et al*, 2007b). In humans and other mammals, four PMCA isoforms (PMCA1-4) are encoded by separate genes (Strehler *et al*, 2007a). PMCA1 is considered a “housekeeping” pump and together with PMCA4 ubiquitously expressed in all tissues. PMCA2 and 3 have specific expression patterns and are mostly found in excitable tissues and often referred to as neuron-specific isoforms (Domi *et al*, 2007; Strehler & Thayer, 2018). Through alternative RNA splicing at two different sites (“A” and “C”), the four isoforms can be produced in more than 20 different variants (Strehler & Zacharias, 2001). Splicing at site C affects the length of the C-terminal tail with CaM-binding sites and splicing at site A impacts the length of the first intracellular loop which leads from the A-domain to the transmembrane segment 3 (TM3) (Strehler, 2015). The different splice variants are in many cases tissue-specific and differ in the degree of activation and autoinhibition (Caride *et al*, 1999; Kessler *et al*, 1992).

PMCA is regulated through interactions with protein partners as well as phospholipids. A classical regulatory feedback mechanism happens via the C-terminal autoinhibitory domain of PMCA. In the resting cell, the domain interacts with the cytoplasmic domains and autoinhibits calcium transport. Upon an increase in intracellular calcium, calcium-bound CaM binds to the autoinhibitory domain resulting in a rise in PMCA activity. The degree and calcium threshold of activation may depend on how many CaM-binding sites – one or two – are present within the autoinhibitory domain (Tidow *et al*, 2012). Furthermore, PMCA regulation by acidic phospholipids has been mapped to two binding sites – one in the autoinhibitory domain, the other at the cytosolic loop between the A-domain and TM3. The mechanism of how acidic lipids regulate the pump is not understood in detail; however, they appear to be both modulators of activation by CaM as well as stand-alone activators (Brodin *et al*, 1992; Lopreiato *et al*, 2014; Niggli *et al*, 1981a; Penniston *et al*, 2014; Pinto Fde & Adamo, 2002; Tidow *et al*., 2012; Zvaritch *et al*, 1990). Hence, the autoregulatory regions also correspond to the sites of variations by alternative splicing.

In the present study, we show that aSN in its soluble, monomeric form acts as a very potent activator of human PMCAs. The effect relies on the presence of acidic phospholipids and is independent of the CaM-binding autoinhibitory domain. Our findings suggest that the activation mechanism is based on interactions of the N-terminal segment of aSN, negatively charged lipids, and a phospholipid binding site of PMCAs. We propose that aSN complements CaM in local compartments such as the presynapse. Our finding gives a new perspective on the physiological role of native aSN in the presynapse and demonstrates how the lipid environment affects the critical calcium extruding activity of PMCA.

## RESULTS

### aSN co-localizes with PMCA and stimulates calcium extrusion

In previous mass spectrometry studies, PMCA was noted as a potential interaction partner of aSN (Betzer *et al*., 2015). Further co-immunoprecipitation experiments analyzed by western blotting show that endogenous PMCA interacts directly or indirectly with endogenous aSN (**Figure 1A, right**). Co-immunoprecipitation experiments with aSN-knockout brain homogenate supplemented with either purified aSN monomer, or *in vitro* formed aSN oligomer revealed that PMCA did not display preferential aSN-conformational binding as interactions occur with both the monomer and the oligomer (**Figure 1A, left**).

**Figure 1.**
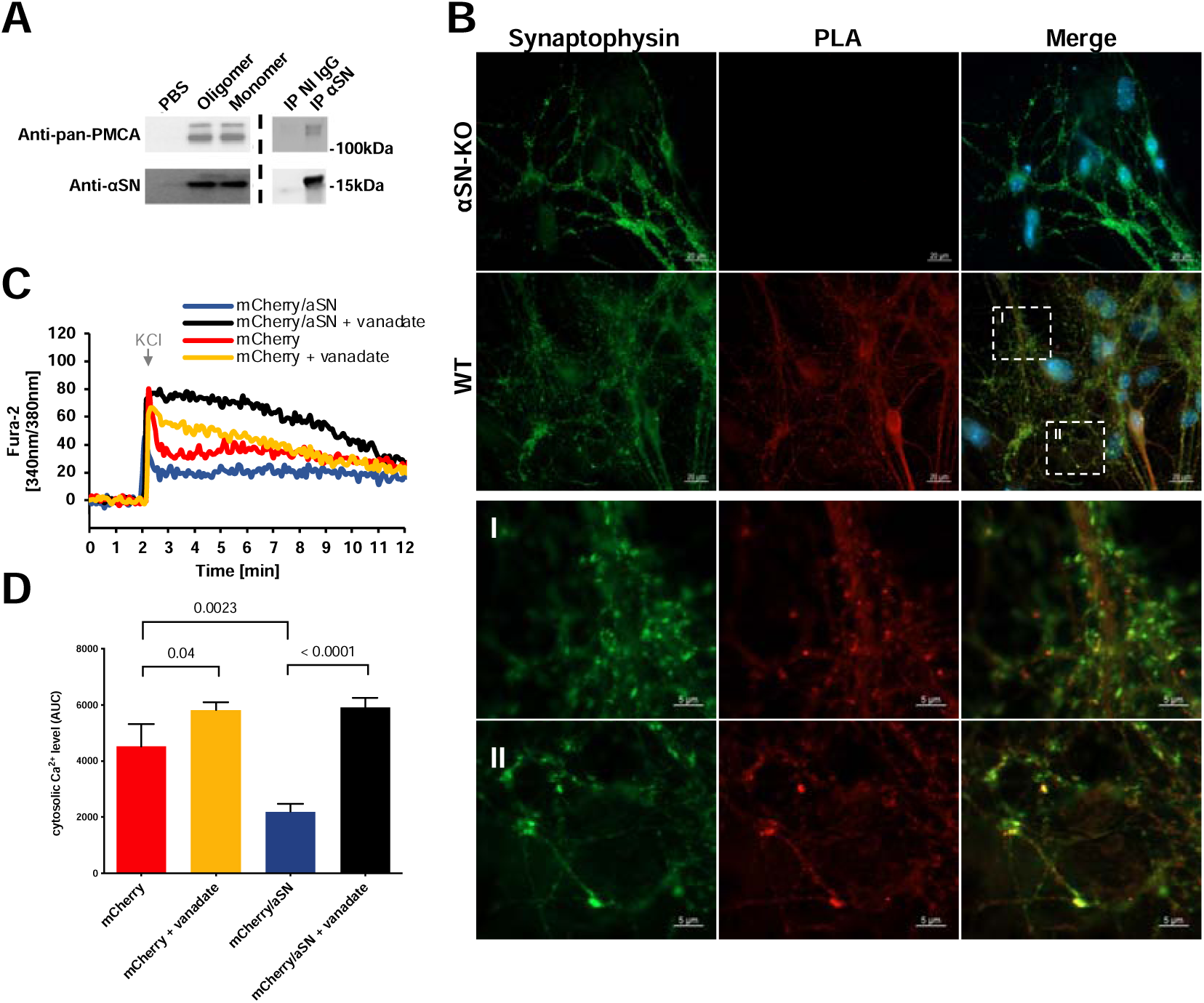
aSN co-localizes with PMCA and stimulates calcium extrusion. **A. Co-immunoprecipitation of PMCA with aSN**. *Left*: In detergent extracts of total brain homogenates from aSN-knockout (aSN-KO) mice both exogenous aSN monomer and in vitro formed oligomers are pulled down together with PMCA mice by aSN binding sepharose (ASY-1). Control without exogeneous aSN (PBS) confirms no unspecific antibody binding. *Right*: In detergent extracts of total brain homogenate from C57BL/6 mice, endogenous aSN is pulled down together with the endogenous PMCA by aSN specific antibody (ASY-1) and not by control non-immune antibody (NI IgG). **B. Proximity ligation assay (PLA) of aSN and PMCA primary hippocampal neurons** PLA of PMCA and aSN in primary hippocampal neurons made from WT or aSN-KO mice. *Left column*: synapses visualized by synaptophysin labeling. *Middle column:* red fluorescence signal represents positive PLA result, meaning proximity of aSN and PMCA of less than 40 nm. *Right column*: Merged PLA and synaptophysin images show PMCA and aSN interaction localized to the synapses. *I and II:* Zoomed-in images of boxed areas of WT primary hippocampal neurons. **C and D. aSN increases calcium export from depolarized primary neurons**. Cytosolic calcium was monitored by Fura-2-AM loaded into DIV8 neurons. Before recording SERCA was inhibited by thapsigargin. Calcium influx was induced by the addition of KCl. At the recording time of 2 min. KCl was added to depolarize the neurons and the calcium response was followed over time. **C** – Representative curves from measurements in: *blue* - neurons expressing mCherry and aSN, *black*: neurons expressing mCherry and aSN treated with 1µM vanadate to inhibit ATPases, *red* – neurons expressing mCherry alone, *yellow* – neurons expressing mCherry alone and treated with 1µM vanadate. **D** – Cytosolic Ca^2+^ level after the KCl-induced influx, quantified as the area under the curve (AUC±SEM). The response upon K^+^ induced depolarization was quantified as the Area Under Curve (AUC) from each measured neuron in the 2-12 min. interval. N (mCherry/aSN) = 7, N (mCherry/aSN + vanadate) =10, N (mCherry) = 6, and N (mCherry + vanadate) =10. The colors of the bars correspond to the top figure. Data presented as mean±SEM. Statistical analysis is conducted as multiple comparisons with one□way ANOVA combined with Sidak post hoc test.

The interaction between aSN and PMCA was investigated further in primary hippocampal neurons isolated from newborn C57Bl/6 mice. Primary hippocampal neurons were analyzed after 14 days in culture by immunofluorescence labeling of synapses by synaptophysin and an aSN/PMCA proximity ligation assay (PLA) yielding a red fluorescent signal when aSN and PMCA are located within 40nm. PMCA and aSN were found to be in this proximity of each other and located to the synapses **(Figure 1B)**.

The functional effect of PMCA calcium transport was investigated in primary hippocampal neurons from aSN-KO neurons transiently transfected with either mCherry oraSN and mCherry. SERCA was inhibited by thapsigargin (4µM) before recording. Cellular calcium responses were monitored by the calcium-sensing dye Fura2-AM upon depolarization by 8 mM KCl in the extracellular medium. Neurons expressing aSN expel calcium to a markedly higher degree than the mCherry expressing neurons **(Figure 1C)**. The calcium expulsion was Ca2+-ATPase dependent as the inhibition by vanadate decreased the calcium expulsion in both the mCherry expressing neurons and the aSN mCherry neurons.

### Monomeric aSN activates human PMCA in a lipid-dependent manner and independent of the autoinhibitory domain of PMCA

The measurements of calcium-dependent ATPase activity of PMCA were performed with two available human PMCA variants for which expression and purification had been established in the laboratory – ubiquitous PMCA1 and neuron-specific PMCA2 isoforms (specifically PMCA1d and PMCA2w/a splice variants). The experiments were performed with PMCAs relipidated either with porcine brain phosphatidylcholine (BPC, neutral lipids) or with bovine brain lipid extract Folch fraction I (from here on referred to as brain extract, BE), 60% of which consist of negatively charged phosphatidylinositol and phosphatidylserine lipids. We observed monomeric aSN causing a strong increase in the activity of both PMCA isoforms, however only for PMCAs relipidated with BE, not BPC. As the titration with Ca^2+^ reveals, the elevation of ATPase activity is accompanied by a great rise in the apparent calcium affinity **(Figure 2A, left, center; Table 1)**. Additionally, the activation is gradually abolished by increasing the content of neutral BPC in the relipidation mixture **(Figure 2A, right)**. To rule out potential, unspecific effects of other ATPases or impurities in the PMCA preparations, we prepared an inactive variant of PMCA (PMCA1d_D475N_), in which the reactive aspartyl residue in the active site was replaced by asparagine. In terms of purity, these preparations were no different from the wild type, as confirmed by SDS-PAGE and size exclusion chromatography **(Figure S1)**, and they showed no ATPase activity, neither with nor without aSN **(Figure 2A left, 2B)**.

**Figure 2.**
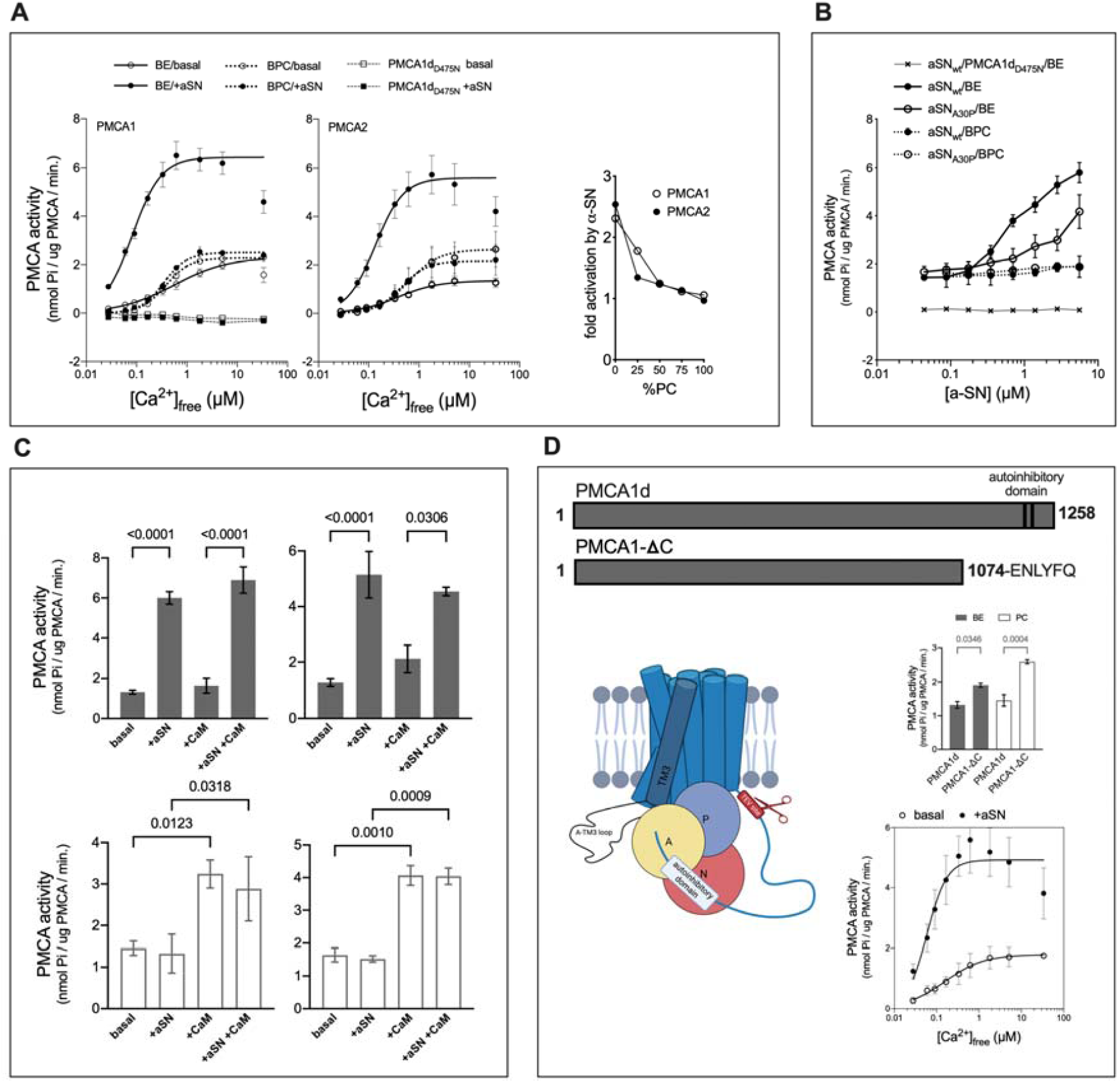
Monomeric alpha-synuclein activates human PMCA in a lipid-dependent and autoinhibitory domain-independent manner. **A. aSN stimulates plasma membrane calcium ATPase activity in an acidic, but not neutral lipid environment**. aSN activates PMCA1d (left) and PMCA2w/a (center) in the presence of brain extract (BE, *solid line*), but not in presence of brain PC (BPC, *dotted line*). PMCA activity was measured as a function of free Ca^2+^ concentration ([Ca^2+^]_free_). *Empty symbols* represent basal activity and *filled symbols* the activity in presence of 2.8 μM aSN. The lines are the best fit given by the Hill equation with *K*_*d*_ values listed in Table 1. To rule out nonspecificity of ATP hydrolysis observed in wild-type PMCA, an inactive D475N mutant of PMCA1d was tested (*squares, dashed line*), Right: Fold activation of PMCAs by aSN decreases with the increasing amount of neutral lipids. Brain PC was titrated in BE. *Empty symbols* – PMCA1d, *filled symbols* – PMCA2w/a. Data in the calcium titration experiment are mean ± s.e.m., for some points, the error bars are smaller than symbols. PC titration data are from a single experiment for each of the PMCA isoforms. **B. Compared to wild-type, A30P aSN has reduced ability to activate PMCA in the presence of acidic lipids**. PMCA1d activity was measured as a function of aSN (wild-type or A30P) concentration in presence of 1.8 μM free Ca^2+^. PMCA1d was relipidated either with BE or BPC. Neither of the aSN variants activated PMCA in a neutral lipid environment. The inactive PMCA1d (D475N) was not stimulated by aSN in the presence of BE. Data are mean ± s.e.m. For some of the points, the error bars are smaller than the symbols. **C. Neutral or acidic lipids create a condition for PMCA being activated exclusively by CaM or a-SN**. Bars display the PMCA1d (*left*) and PMCA2w/a (*right*) activity without activator, with aSN, with CaM, or both. PMCA was relipidated in BE (*top, filled bars*) or BPC (*bottom, empty bars*). **D. The activation by aSN is independent of PMCA’s autoinhibitory domain**. *Top*: Comparison of two PMCA1 variants – full-length PMCA1d with CaM-binding sites and PMCA1-ΔC, where the C-terminal tail, containing the autoinhibitory domain with CaM-binding sites, was removed by cleavage with TEV protease. TEV proteolytic cleavage site was introduced by site-directed mutagenesis at residue 1074. *Bottom left*: Schematic representation of PMCA1d with TEV cleavage site introduced by site-directed mutagenesis. *Middle right*: Basal activity of PMCA1d and PMCA1-ΔC in presence of either BE or BPC. *Bottom right*: The activity of PMCA1-ΔC with (*filled symbols*) and without 2.8 μM aSN (*empty symbols*) as a function of increasing [Ca^2+^]_free_. PMCA1-ΔC was relipidated with BE. The lines are the best fit given by the Hill equation with *K*_*d*_ values listed in Table 1. Except for cases of Ca^2+^ or aSN titration, the fixed concentrations were 1.8 μM or 5.6 μM, respectively. Relevant no-calcium or no-aSN backgrounds were subtracted. Data from at least three independent experiments are expressed as mean ± s.e.m. Statistical analysis was conducted in GraphPad Prism software as multiple comparisons with one□way ANOVA combined with Sidak post hoc test. Subfigure 2B (*bottom left*) as well as 3C, 3D (*bottom right*), 4A, 4C and 4E were created with Biorender.com

**Table 1.**
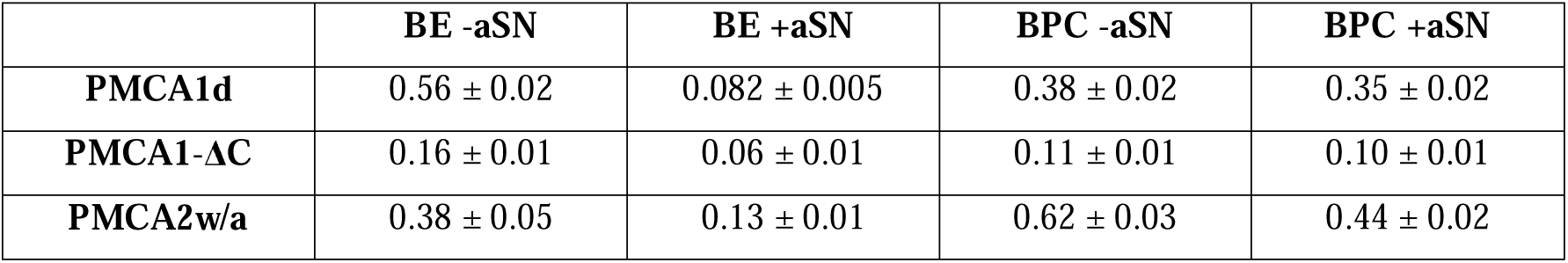
*K*_*d*_(Ca^2+^) values (μM) of PMCAs in different lipid environments. BE – brain lipid extract containing negatively charged phosphatidylinositol and phosphatidylserine; BPC – brain phosphatidylcholine (PC). Value ± SEM given by the Hill equation.

|  | BE -aSN | BE +aSN | BPC -aSN | BPC +aSN |
| --- | --- | --- | --- | --- |
| **PMCA1d** | 0.56 ± 0.02 | 0.082 ± 0.005 | 0.38 ± 0.02 | 0.35 ± 0.02 |
| **PMCA1-ΔC** | 0.16 ± 0.01 | 0.06 ± 0.01 | 0.11 ± 0.01 | 0.10 ± 0.01 |
| **PMCA2w/a** | 0.38 ± 0.05 | 0.13 ± 0.01 | 0.62 ± 0.03 | 0.44 ± 0.02 |

To further investigate the acidic phospholipid dependence, we performed aSN titration comparing effects of wild type aSN and its A30P variant with reduced membrane-binding ability (Jensen *et* al., 1998). In BE, the activation of PMCA was significantly attenuated by the A30P mutation. As could be expected, the aSN variant was ineffective in the neutral environment of BPC (**Figure 2B**).

Next, we examined the potential interplay between aSN and the best-known protein activator of PMCA – CaM. For both PMCA isoforms, the susceptibility for activation by CaM and aSN appears to be lipid-dependent, with the acidic lipids promoting the effect of aSN, and neutral lipids promoting the effect of CaM **(Figure 2C)**. To confirm that the action of aSN was independent of the autoinhibitory domain, we designed a truncated construct (PMCA1-ΔC), lacking 184 residues of the C-terminus including the autoinhibitory domain with the CaM-binding site and a long, intrinsically disordered linker region **(Figure 2D, top)**. Foreseeably, loss of the C-terminal tail resulted in a constitutively active protein, insensitive to CaM **(Figure S3)** with basal activity higher than that of the full-length wild type (**Figure 2D, bottom left)**. However, the stimulation by aSN was preserved and appeared similar to the full-length pump, indicating the aSN effect to be independent of the autoinhibitory mechanism **(Figure 2D, bottom right)**. The apparent Ca^2+^ affinity of PMCA1d was increased by the truncation itself, and aSN strengthened that effect **(Table 1)**. Again, the activation was lipid-dependent, occurring in the PMCA1-ΔC variant reconstituted in BE and not in BPC **(Figure S3)**.

Additionally, we examined the oligomeric form of the full-length aSN, which was previously established to have a great activating impact on the SERCA (Betzer *et al*., 2018). For PMCA, however, the effect was smaller than that of the monomer, occurring only at higher concentrations and to a lower fold of activation **(Figure S3D)**. The activation might simply be explained by dissociating monomer, which we have found to be present in oligomer preparations **(Figure S2)**.

### PMCA alternative splicing events correlate with a-SN expression level

PMCA is extensively regulated by alternative splicing, which results in over 20 variants, differing in distribution, kinetic properties, and sensitivity to interaction partners e.g., CaM (Krebs, 2015; Strehler & Zacharias, 2001). We took into consideration that some of them could vary also in how they respond to aSN. We used available transcriptomics data from the VastDB database (http://vastdb.crg.eu/), containing information on alternative splicing events. The level of alternative splicing events in different tissues are indicated by PSI values (“percent spliced in”), and we asked whether specific splicing events correlate with aSN expression levels. First, we analyzed the splice site C, which is located at the C-terminal region of PMCA and contains the autoinhibitory domain that interacts with CaM. Alternative splicing at this site gives variants that may differ in the degree of autoinhibition or rate of activation by CaM. The splice variant “a”, with a shortened autoinhibitory domain is known to have higher basal activity and to be less sensitive to CaM stimulation (Caride *et al*, 2007). The available data regarded PMCA1, PMCA3, and PMCA4, and the analysis displayed in **Figure 3A** shows that in the case of PMCA1 and PMCA4 exon, incorporation leading to the variant “a” positively correlates with the expression level of aSN with only a few outliers. Only PMCA3 did not correlate and the PSI value of the “a” variant was relatively high regardless of aSN expression interval. The results suggest that in tissues highly expressing aSN, PMCA is alternatively spliced in a way that favors less CaM-dependent variants. The PMCA “a” variant is mainly expressed in brain tissues and explains the positive correlation between PMCA and aSN. Except for brain tissues, aSN is also expressed in high amounts in melanocytes and bone marrow, which account for the two outliers in the plot.

**Figure 3.**
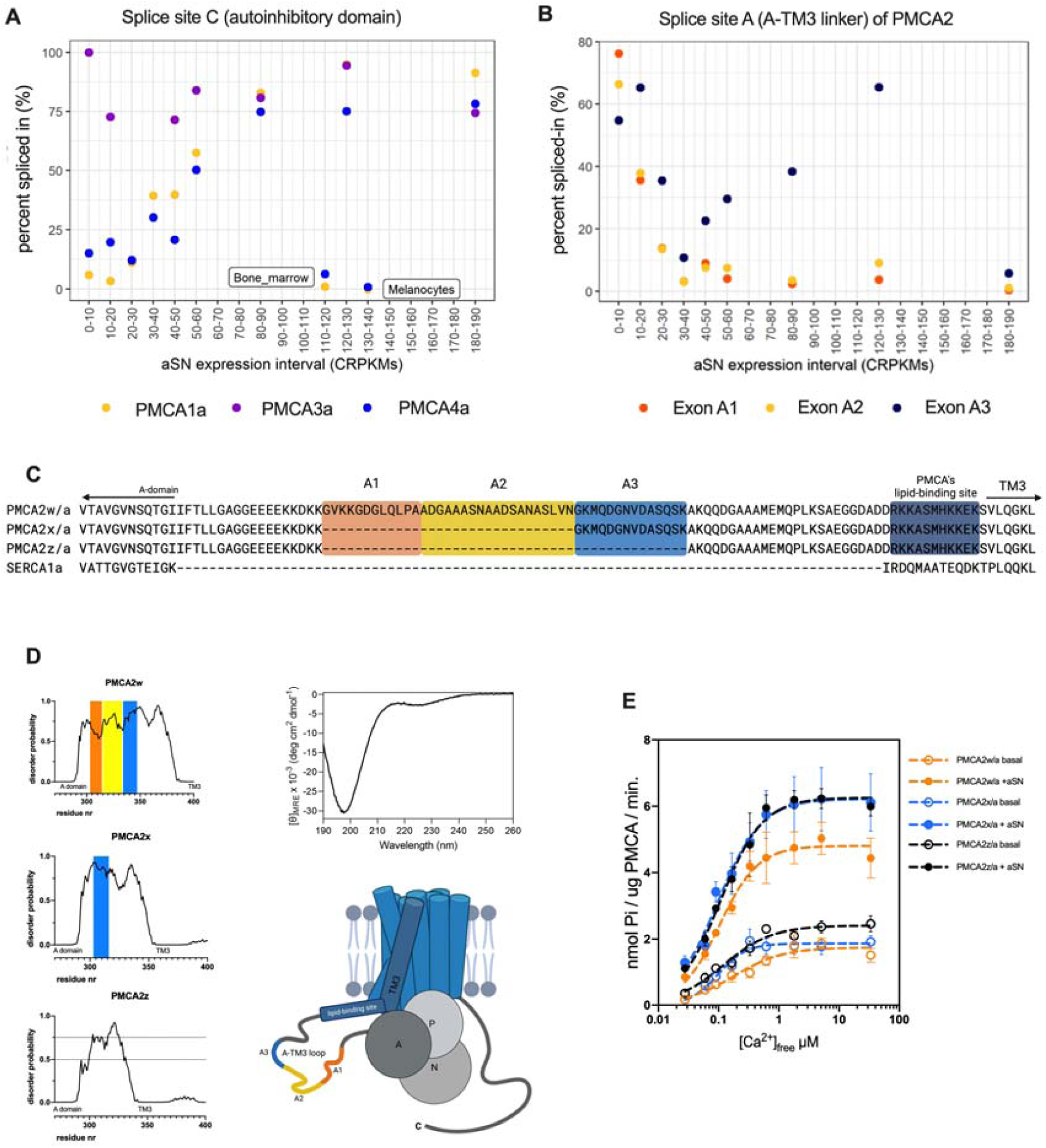
Alternative splicing events of PMCA correlate with aSN expression levels and can modulate activating effect of aSN. **A-B. PMCA’s alternative splicing events correlate with a-SN expression levels**. The analysis of transcriptomics data from the VastDB database. A – Exon incorporation for exons at splice site C for PMCA 1, 3, and 4. The average value of percent spliced in (PSI) for the exon leading to splice variant “a” as a function of aSN tissue expression. B – Exon incorporation for exons at splice site A for PMCA 2. The average value of PSI for the three possible exons as a function of aSN tissue expression. The color code for the exons (orange, yellow, blue) is consistent with corresponding PMCA regions marked on the subfigures C and D. For VastDB entry numbers of analyzed events refer to the method section **C. Sequence alignment of the extended A-TM3 loop of PMCA2w, x, and z variants**. Fragments A1, A2, and A3 of the A-TM3 loop correspond to alternative exons incorporations. Color code corresponds to respective alternative splicing events presented in figure B. Sequence of the corresponding region of SERCA1a for comparison. **D. Structural disorder in the A-TM3 loop of PMCA2**. Disorder predictions: *Top left* – PMCA2w, *middle left* – PMCA2x, *bottom left* – PMCA2z, and *top right*, far-UV CD analysis of PMCA2w_A-TM3_ loop with minimum below 200 nm reflecting a high degree of disorder. *Bottom right* – schematic representation of PMCA2w/a with the intrinsically disordered loop region between domain A and TM3. **E. Comparison of aSN impact on three splice variants of human PMCA2** – Ca^2+^-dependent activity of PMCA2w, x and z. The lines are the best fit given by the Hill equation with the apparent *K*_*d*_ values (μM) for Ca^2+^ as follows: PMCA2w/a basal (without aSN) – 0.18 ± 0.04, with aSN 0.102 ± 0.009; PMCA2x/a basal – 0.10 ± 0.01, with aSN 0.097 ± 0.009; PMCA2z/a basal 0.13 ± 0.02, with aSN 0.106 ± 0.008. Values calculated in GraphPad Prism

The other of the two splice sites – A – is located at the lengthy A-TM3 linker region between the A-domain and TM3 (from here on referred to as “A-TM3 loop”). Among isoforms, PMCA1 is not being spliced in this region, while in PMCA2 alternative exon composition leads to 4 different variants named “z’, “x”, “y”, and “w”, with all of them except “y” having been detected in humans (Di Leva *et al*, 2008; Strehler *et al*., 2007b). **Figure 3B** displays the analysis of the PMCA2 splicing events, where three exons can be incorporated leading to variants “w” (full length containing sequences marked in orange, yellow and blue) and “x” (containing the blue sequence). The “z” variant occurs when none of the three exons are incorporated **(Figure 3C)**. We observed incorporation of the exons leading to variants with a longer A-TM3 loop insert in tissues with low aSN expression and barely any incorporation of these exons for tissue with high expression of aSN. This suggests that low expressing tissues would preferentially transcribe the longest PMCA2w, and tissues richer in aSN would express more PMCA2x and z.

The extended, variable A-TM3 loop of PMCA is predicted to be structurally disordered; a trait which was confirmed using circular dichroism (CD) spectroscopy analysis of a recombinantly expressed PMCA2w_A-TM3_ loop **(Figure 3D)**. Near TM3, the loop contains several positively charged residues that mediate the activation by acidic phospholipids (Brini *et al*, 2010; Brodin *et al*., 1992; Pinto Fde & Adamo, 2002). We speculated about the importance of the loop length in the interaction with disordered aSN and performed the free calcium titration experiment with PMCA2 variants w/a, x/a, and z/a. All PMCA2 constructs used in the experiment were transformed anew into yeast cells derived from one colony and were expressed, purified, and tested simultaneously. The activity assays show that all three variants are activated by aSN **(Figure 3E)**. The most remarkable difference is that PMCA2w/a has the lowest maximum activity among variants when activated by aSN. It was also the only one with a substantial increase in the apparent calcium affinity induced by aSN, the other having high affinity also in the basal activity state. PMCA2z/a, having the shortest loop region, showed the highest basal activity. The results implicate the A-TM3 loop in aSN interactions.

### Indirect interaction between aSN and PMCA occurs between the N-terminal region of aSN and the acidic phospholipid binding site of PMCA

To determine whether the activation of PMCA was triggered by a direct interaction of aSN with the PMCA A-TM3 loop, we performed interaction studies using NMR spectroscopy. We incubated recombinant PMCA2w_A-TM3_ loop with ^15^N-labelled aSN and found that it induced no chemical shift perturbations (CSPs; **Figure S4A**), in both the absence (**Figure S4B**), or presence of Ca^2+^ (**Figure S4C**). Further, we found no indications of a direct interaction between full-length PMCA2w/a and aSN as no CSPs (**Figure S4D**) or changes in NMR peak intensities (**Figure S4E**) in the aSN spectra could be seen. This suggests the interaction between aSN and PMCA is dependent on a more complex environment and likely is occurring via a mechanism facilitated by the membrane.

To map the potential interaction site on the aSN sequence (1-140) we performed a comparison of shortened aSN variants, truncated N-terminally (Δ1-14, Δ1-29) to compromise membrane binding (Cholak *et al*, 2020) or C-terminally (1-95, 1-61) to compromise calcium binding and to remove the NAC binding region (Skaanning *et al*, 2020) respectively **(Figure 4AB)**. Most notably, the Δ1-14 variant had a greatly weakened effect on PMCA and Δ1-29 had almost lost its ability to activate PMCA, displaying only minor effects above 5 μM concentration. The C-terminally truncated (1-95) variant generally saved function although to a lesser extent, activating to a lower fold (Figure 4B, Supplementary figure 3B). The (1-61) variant of a-SN did not reach saturation within the examined concentration range, which suggests greatly reduced apparent affinity to PMCA.

**Figure 4.**
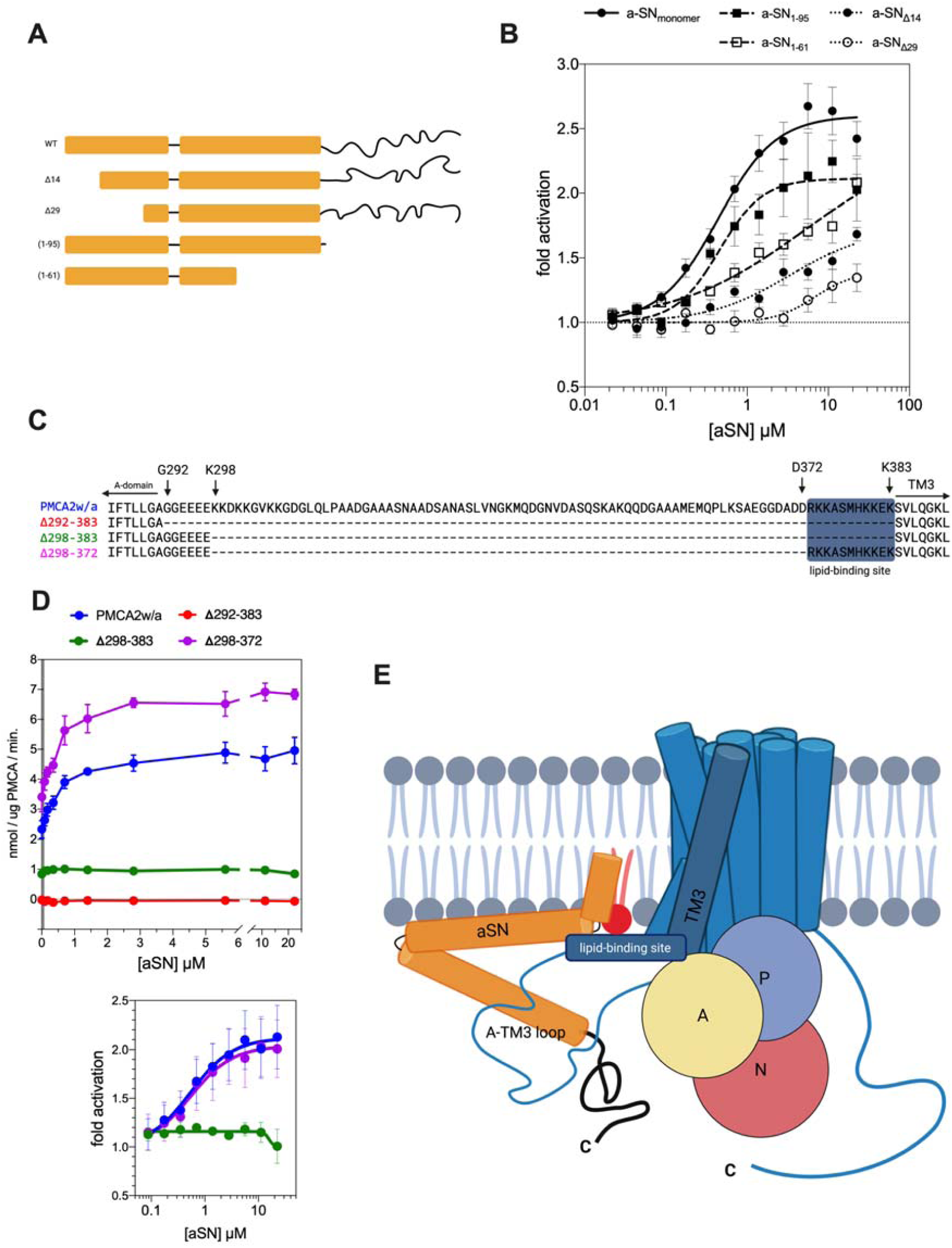
The aSN-PMCA interaction involves the N-terminal region of aSN and the acidic phospholipid binding site of PMCA mediated by acidic phospholipids. **A. Schemes of aSN and its N- and C-terminal truncation variants**. Bars represent facultatively (lipid-mediated) helical regions, lines – constitutively disordered regions. **B. Membrane anchoring of aSN is important for the activation of PMCA**. PMCA1d fold activation by titrated full-length aSN and truncated variants. The experiment was performed in presence of 1.8 μM free Ca^2+^ and brain lipid extract was used for the PMCA relipidation. Data are a result of at least three independent activity assays; mean ± SEM. **C.** Sequence comparison of full-length PMCA2w/a and its deletion variants, missing large fragments of the A-TM3 loop: Δ292-383, Δ298-383 Δ298-372. The blue background indicates the acidic phospholipid-binding site. **D. The PMCA’s acidic phospholipid binding site preceding the TM3 is crucial for the activation by the aSN**. *Top* – The specific activity of PMCA in the presence of 1.8 μM free Ca^2+^. PMCA2w/a and variants were relipidated in the BE and the full-length monomeric aSN was used as the activator. *Bottom* – baseline-corrected data displayed as fold activation of PMCA by aSN. Data are the result of at least three independent measurements; mean ± SEM. **E. Proposed mechanism of the PMCA-aSN interaction**. In the presence of acidic membrane phospholipids (*red*) the N-terminal region of aSN anchors it to the plasma membrane *vis-a-vis* acidic lipids, which mediate the interaction with the acidic lipid-binding site of PMCA, leading to activation of PMCA.

The disordered C-terminal tail of aSN has calcium-binding properties (Lautenschlager *et al*., 2018) with a relevant affinity of ∼20-50 *μ*M (Newcombe *et al*, 2021) thus we asked if it would impact the apparent calcium affinity of PMCA. For this, we compared activation of PMCA by the full length and C-terminally truncated (1-95) aSN in the Ca^2+^ titration experiment. However, despite the lower fold-activation, the C-truncated construct did not differ in shifting PMCA’s apparent *K*_*d*_ for Ca^2+^ **(Figure S3B)**.

Within certain limitations, deletions of large fragments of the lengthy A-TM3 loop of PMCA, including its acidic lipid-binding site, can retain basal activity (Pinto Fde & Adamo, 2002). Guided by this, we designed constructs and expressed and purified certain deletion variants of PMCA2, where we removed large parts of the loop. Sequences of the targeted region of PMCA2w/a and the three designed variants (Δ292-383, Δ298-383, Δ298-372) are displayed in **Figure 4C**. The activity assay in the presence of BE **(Figure 4D)** showed differences in the basal activity and response to aSN. We observed a complete loss of ATPase activity of the Δ292-383 variant. Keeping the six N-terminally located residues of the loop resulted in the Δ298-383 variant with the basal activity preserved (0.84 ± 0.02 nmol Pi/μg PMCA/min.), but not responding to aSN titration. Preserving further the 11-residues long binding site for negatively charged lipids at the C-terminal end of the loop, close to TM3 (Δ298-372) boosted the basal activity (3.4±0.5 nmol Pi/μg PMCA/min.) and reinstalled the fold-activation by aSN to the level observed in wild type PMCA2w/a (but having lower basal activity of 2.3±0.3 nmol Pi/μg PMCA/min). As this points to the binding site for negatively charged lipids as crucial for the interaction with aSN, we propose the aSN-PMCA interaction to be mediated by acidic phospholipids involving the N-terminal region (1-95) of aSN and the acidic lipid binding site of PMCA **(Figure 4E)**.

### The activation by aSN is specific to mammalian calcium pump

PMCA activation by aSN is observed both for the ubiquitous PMCA1 and the more tissue-specific PMCA2 and was also verified for the C-terminally truncated PMCA4x isoform (**Figure S3C**), which likely colocalizes with aSN in synaptic terminals of retinal neurons (Bodis-Wollner *et al*, 2014; Cali *et al*, 2017; Krizaj *et al*, 2002). The broad impact on PMCA isoforms raised the question of whether the stimulatory effect of aSN is at all specific to PMCA.

Interestingly, we found that rabbit SERCA1a is also activated by the aSN monomer in the same lipid-dependent manner as PMCA **(Figure 5A)**. However, the effect was statistically relevant only in low (60 nM) free Ca^2+^ concentration, whereas at high 1.8 μM free Ca^2+^, SERCA was activated to a lower fold. This indicates that aSN may also modulate the apparent Ca^2+^ affinity of SERCA. With continuous ER extending into presynaptic compartments (Singh *et al*, 2021; Wu *et al*, 2017) this is also of relevance to aSN function, and most likely it is also related to the previously reported activation by aSN oligomers (Betzer *et al*., 2018).

**Figure 5.**
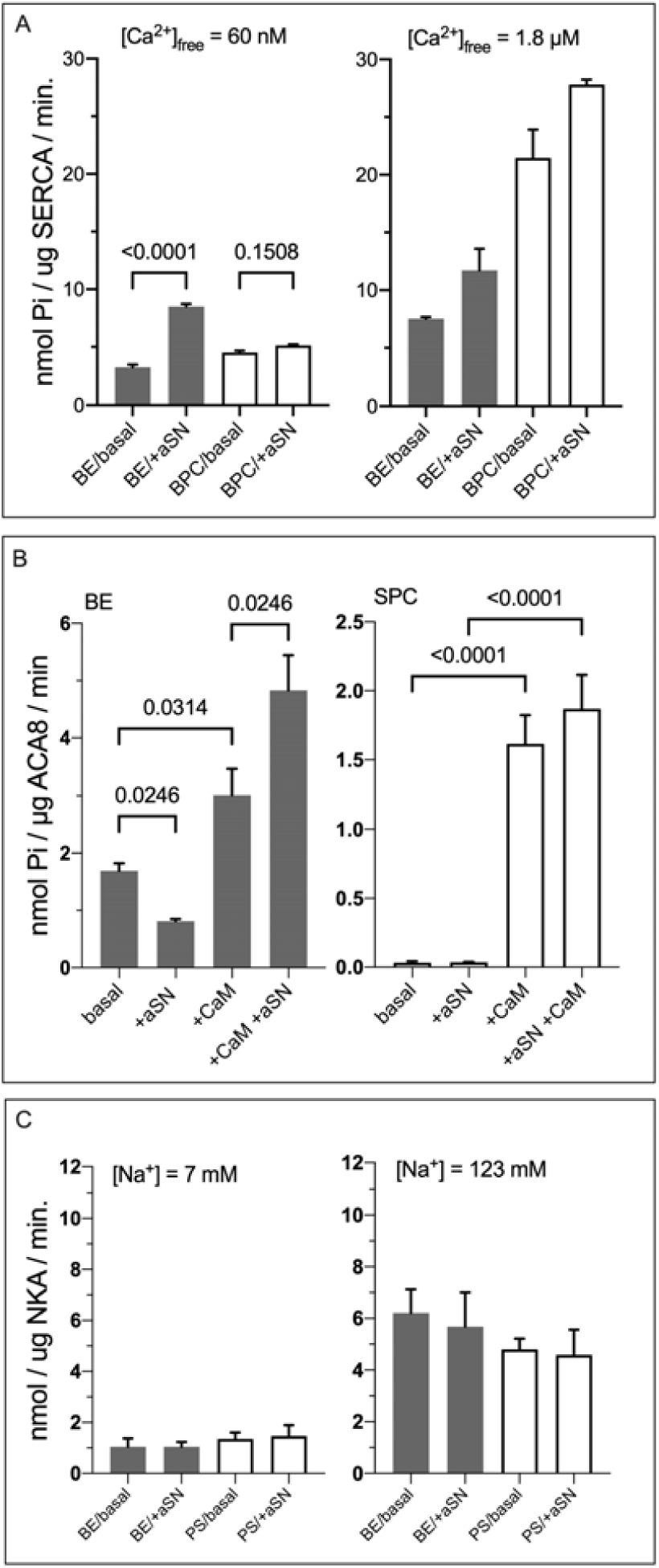
Activation by aSN appears specific to mammalian calcium pumps. **A. In presence of acidic lipids aSN activates rabbit SERCA1a at low [Ca**^**2+**^]_**free**_ **(60 nM)**. Bars show SERCA activity without or with aSN when the pump is relipidated in BE (*filled bars*) or brain BPC (*empty bars*). [Ca^2+^]_free_ is 60 nM (*on the left*) and 1.8 μM (*on the right*). **B. Plant Ca**^**2+**^ **ATPase, ACA8 is not activated by aSN**. Bars show the activity of ACA8 in presence of 1.8 μM Ca^2+^ and conditions: without protein partners, with aSN, with CaM, or with both. PMCA was relipidated in BE (*left, filled bars*) or soy PC (SPC) (*right, empty bars*). **C. Human Na.K ATPase (NKA) is not activated by aSN**. Bars show the activity of NKA in the presence of 7 or 63 nM Na^+^ (right and left graphs, respectively) without or with aSN. NKA was relipidated either in phosphatidylserine (PS) or brain extract (BE). When aSN or CaM are present, the concentrations are 5.6 μM or 1.2 μM, respectively. No-substrate background values were subtracted. Data from at least three independent experiments are expressed as mean ± SEM. Statistical analysis is conducted as multiple comparisons with one□way ANOVA combined with Sidak post hoc test.

Next, we turned to a plant homolog of PMCA, namely the autoinhibited calcium ATPase of *Arabidopsis thaliana* (ACA8), belonging to the same P2B subfamily of P-type ATPase (Axelsen & Palmgren, 2001), but not coexisting with aSN, which is only found in vertebrates. Structurally, ACA8 and PMCA differ in the localization of the autoinhibitory domain, which is situated on the N-terminus of ACA8 and the C-terminus in PMCAs, and ACA8 is lacking the long, disordered loop for the A-TM3 loop with the putative lipid-binding site in PMCA (Brini *et al*., 2010; Brodin *et al*., 1992; Pinto Fde & Adamo, 2002). In the presence of BE, ACA8 is not activated by aSN, but rather inhibited, and the effect is partially reversed by CaM **(Figure 5B)**. In the presence of soy PC, the pump is autoinhibited and can be activated by CaM, but not aSN. Furthermore, the human Na.K-ATPase (*α*1 isoform) was not activated by aSN **(Figure 5C)**. These results suggest the specificity of the aSN interaction for mammalian Ca^2+^-ATPases.

## DISCUSSION

Here we show a strong, activating effect of soluble, monomeric aSN on PMCAs and to a certain degree also SERCA. Our results suggest that the physiological role of aSN is stimulation of calcium clearance by increasing V_max_ and/or apparent calcium affinity of the active transporters. Due to the very specific localization of aSN, this mechanism would be restricted to specialized cells or cellular compartments like presynaptic termini of neurons. This is supported by our proximity ligation study showing an intracellular colocalization of PMCA and aSN in nerve terminals, and by experiments in primary neurons where we show that aSN increases calcium expulsion after depolarization.

The charge of phospholipids in the PMCA environment has a great influence on its regulation, and acidic lipids alone can activate PMCA to some extent via specific interaction sites (Carafoli & Stauffer, 1994; Lopreiato *et al*., 2014; Strehler, 2015). In a similar way to the recently found inhibition by tau protein (Berrocal *et al*, 2017), we found the effect of aSN on PMCA is enabled by negatively charged and not neutral phospholipids in the membrane. The opposite lipid dependence is observed for CaM, consistent with earlier findings (Lopreiato *et al*., 2014; Niggli *et al*., 1981a; Niggli *et al*, 1981b). The composition of the local lipid environment could potentially switch PMCA modes from being responsive either to aSN or CaM.

With its strong effect and accumulation at presynaptic compartments, aSN would allow for a rapid, stimulated response to incoming calcium signals and integration with other calcium-sensitive presynaptic functions such as neurotransmitter release. This, together with aSN activating independently of the autoinhibitory C-terminal tail of PMCA, suggests aSN complements CaM as the activator of PMCA in this specific neuronal compartment.

The alternative splicing of PMCA generates many variants among isoforms and has been for a long time proposed to be a mechanism of fine-tuning of calcium handling in cells affecting expression, localization, and membrane trafficking to specific compartments (Antalffy *et al*, 2011; Krebs, 2015; Strehler & Zacharias, 2001). Splice site “A” variants of PMCA (w,x,y,z) differ in the size of the lengthy and disordered A-TM3 loop (**Figure 3C**). That region was not resolved in a cryo-EM structure of PMCA1 (Gong *et al*, 2018), and its impact on the enzymatic properties is not clear, but the proximity to the membrane interface and cytosolic Ca^2+^ entry suggests a modulatory role of the A-TM3 loop; also its involvement in PMCA-lipid interaction and membrane trafficking has been proposed (Adamo & Penniston, 1992; Antalffy *et al*., 2011; Enyedi & Strehler, 2011). Our mutational studies indeed indicate the modulation of the activity. In PMCA, the deletion of the intrinsically disordered A-TM3 loop (deleting residues 298-383 in PMCA2w/a) retained basal enzymatic function, but the protein was unresponsive to aSN. However, the stimulation was preserved in a Δ298-372 variant, where 11 residues upstream of the TM3 are not deleted. This fragment, highly specific in sequence to PMCA, is rich in positive charge and was previously shown to interact with acidic lipids (Adamo & Penniston, 1992; Pinto Fde & Adamo, 2002). Additionally, this variant showed increased basal activity, which may indicate a role of the A-TM3 loop in some level of autoinhibition that complements the autoinhibitory domain at the C-terminal end of PMCA.

The correlations found between aSN expression level and splicing events at the “C” site of PMCA isoforms 1 and 4 suggest that an “aSN/CaM” balance of PMCA activation can be managed by alternative splicing, leading to the synthesis of less CaM-sensitive variants in tissues destined for aSN activity. Variations obtained at the splice site “A” of PMCA2 suggest preference for variant “w” in low-aSN-expressing tissues and shorter variants “x” and “z” being expressed together with increasing aSN level. In the activity assay, the PMCA2 splice variants responded differently to aSN: PMCA2w/a was the only one having its apparent calcium affinity increased by aSN, but on the other hand, it was stimulated to the lowest maximum activity and had relatively low apparent calcium affinity in a basal (non-activated) state. The longest A-TM3 loop of PMCA2w can through its capacity of its long disorder (**Figure 2D**) be associated with more flexibility in the regulation of its affinity and capacity. In PMCA2”x” and “z” only the capacity was stimulated. The last two variants would be expressed in aSN-abundant tissues with potential demand for higher capacity for calcium extrusion.

The specific function of soluble, monomeric aSN is debated (Cheng *et al*, 2011; Sulzer & Edwards, 2019), but preferential association to membranes with acidic phospholipids is a recurring theme (Davidson *et al*, 1998; Jo *et al*., 2000). The ability to activate PMCA appears strongly related to its lipid-binding properties. The aSN A30P mutant form, associated with familial early-onset PD, is known for reduced ability to bind acidic phospholipids (Jo *et al*, 2002), and that correlates with the weaker PMCA activation. Similar dependence emerges from the characteristics of the N-terminus of aSN. Monomeric aSN interacts with acidic phospholipids via the ∼93 residues long N-terminal part adopting amphipathic helical segments (Ulmer *et al*, 2005; Uversky & Eliezer, 2009). The aSN binding to lipid bilayers has been proposed to be maintained by avidity between the 1-14 region, strongly interacting and anchoring into the lipid bilayer, and the downstream amphipathic helix having weak surface binding capacity (Cholak *et al*., 2020). Indeed, here we observed a gradual loss of the PMCA activating properties by deletion of 1-14 or 1-29 residues, with the Δ(1-29) construct being almost incapable of activating PMCA.

From a structural and mechanistic point of view, we propose that PMCA is activated by through a *vis-à-vis* interaction between the N-terminus of the monomeric aSN and the acidic lipid binding site of PMCA, located at the C-terminal side of the disordered A-TM3 loop, and mediated by acidic lipids. The bulk part of the loop can play a tuning role together with the disordered C-terminal tail of aSN. With low-affinity Ca^2+^-binding properties (K_d_ ∼20-50 μM) (Lautenschlager *et al*., 2018; Newcombe *et al*., 2021), this region of aSN could potentially increase the local concentration of Ca^2+^ at the cytosolic entry site of PMCA. It appears to play a role in PMCA activation capacity-wise but does not have a visible impact on apparent calcium affinity.

The lipid-binding site of PMCA is likely functionalized by the many positively charged residues in the 11-residue sequence defining it (RKKASMHKKEK, see also **Figure S5**). The equivalent sequence in the plant orthologue ACA8 (MASISEDNGEE, residue 352-362) is negatively charged, as is the N-terminal part of the A-TM3 loop in the PMCA2-Δ298-383 (TLLGAGGEEEE). Both show significant basal activity in presence of acidic lipids but are not activated by aSN. The corresponding region in SERCA (RDQMAATEQDK, residue 236 to 246 of rabbit SERCA1a) shows a mixed positive and negative charge, which may explain a mild activation by aSN and negatively charged lipids. We note that aSN contains several K-rich motifs in its N-terminal region (1-61) that have resemblance to the acidic-lipid binding sequence of the disordered loop of PMCA; an observation that together with the absence of a direct interaction in simple systems (**Figure S4**) and a neurotoxicity effect of their mutation (Dettmer *et al*, 2015), strengthen the suggestion of a lipid-mediated interaction as the mechanism of aSN activation of PMCA in neurons.

The function of aSN oligomers and fibrils has been thoroughly studied and among a plethora of cytotoxic capacities they have been linked to calcium dysregulation (Betzer *et al*., 2018; Danzer *et al*, 2007; Rcom-H’cheo-Gauthier *et al*., 2016). We show here that native, monomeric aSN exerts a strongly activating effect on PMCA, whereas oligomeric aSN was weaker and perhaps even just ascribed to the small amount of released monomer **(Figure S2)**.

This novel function of aSN is relevant to calcium homeostasis of neurons and specifically presynaptic compartments; however, potentially affects also extracellular environment, where the exchange of each calcium ion for two protons by PMCA leads to formation of transient alkaline nanodomains of the synaptic cleft with consequences to postsynaptic NMDAR fluxes (Chen & Chesler, 2015; Feghhi *et al*, 2021).

The role of aSN in hematopoietic system has been discussed recently (Pei & Maitta, 2019). The interaction with PMCA can be potentially important to calcium especially in red blood cells, where aSN is very abundant (Barbour *et al*, 2008). Erythrocytes lack ER and mitochondria, and their calcium homeostasis depends solely on PMCA. The affinity-increasing effect of abundant aSN on PMCA could explain a very low resting calcium level at 30-60 nM observed in red blood cells (Bogdanova *et al*, 2013).

In the presynapse, PMCA-dependent control of the local calcium-depletion zones could be enforced by aSN and acidic lipids. This may for instance have importance for synaptic vesicle release and recycling where PMCA was proposed to functionally separate simultaneous calcium signals (Krick *et al*, 2021). Oligomerization and aggregation/fibrillation of aSN on the other hand would impair this function and lead to impaired calcium handling. Our finding can then also have implications regarding aSN pathology. Lipid-related dysfunctions have been linked to PD (Fanning *et al*, 2020). Early manifestations of PD could result from calcium dyshomeostasis caused by loss of function by aggregating aSN or lipid-affinity affecting mutations like A30P, which leads to early neuronal dysfunctions (Kruger *et al*, 2001), or changes in the distribution and content of acidic plasma membrane lipids.

## ACKNOWLEDGEMENTS

We thank Torben Heick Jensen and Thomas Gonatopoulos-Pournatzis for advice on transcriptomics data analysis. We are grateful for biosamples and valuable discussion to Michael Habeck for Na,K-ATPase, Thomas Lykke-Møller Sørensen, Anne Lillevang and Claus Olesen for SERCA, Annette E. Langkilde for aSN Δ14 and 1-61. We thank Joseph A. Lyons and Magnus Kjærgaard for valuable discussions on membrane proteins and intrinsically disordered proteins, and Robert Edwards for valuable discussion on alpha-synuclein. We are grateful to Anna Marie Nielsen, Tetyana Klymchuk and Jacob H. Martinsen for technical assistance.

This work was supported by a “Mobility plus, 3^rd^ edition” fellowship grant of the Polish Ministry of Science and Higher Education to A.K., a postdoctoral research grant from the Extremadura Province (PO17009) to M.C.B., by a PhD stipend from the Aarhus Graduate School of Science at Aarhus University to S.T.L., by a collaborative grant from H. Lundbeck A/S to P.H.J., by Lundbeck Foundation grants R223-2015-4222 for P.H.J. and R248-2016-2518 for Danish Research Institute of Translational Neuroscience-DANDRITE, Nordic-EMBL Partnership for Molecular to P.N and P.H.J., and by funds from a project 2 research grant from the Independent Research Fund Denmark and the Brainstruc research center and a professorship grant of the Lundbeck Foundation to P.N. Further support was given by the European Union’s Horizon 2020 research and innovation programme under the Marie Sklodowska-Curie grant agreement No. 101023654, awarded to E.A.N. and the Novo Nordisk Foundation (#NNF18OC0033926 to B.B.K). All NMR data were recorded at cOpenNMR, an infrastructure facility funded by the Novo Nordisk Foundation (#NNF18OC0032996).

## AUTHOR CONTRIBUTIONS

P.H.J. and P.N conceived PMCA studies, P.H.J. and A.K. the aSN interaction studies and E.A.N and B.B.K the in vitro studies with the A-TM3 loop; A.K designed constructs and experiments, except co-immunoprecipitation, proximity ligation assay, and calcium measurements, which were designed, performed and analyzed by C.B. and E.G.; A.K. developed PMCA expression and purification protocols, assisted also by S.T.L. and M.C.B.; C.B. performed the expression and purification of aSN; A.K., S.T.L., and M.C.B. performed and analyzed ATPase activity assays; S.T.L. expressed and purified ACA8 and CaM, and performed the analysis of the transcriptomics data; E.A.N. purified proteins and performed all NMR and CD experiments; A.K. wrote the manuscript with P.N. and with inputs from all authors.

## METHODS

### RESOURCE AVAILABILITY

#### Lead contact

Requests for further information, resources, and reagents should be directed to and will be fulfilled by the lead contact, Poul Nissen

#### Materials availability

All plasmids generated in this study are available from the Lead Contact without restriction. Reagents used in the study were of general use and from commercial sources.

#### Data and code availability

The transcriptomics data are publicly available at https://vastdb.crg.eu/ All original code written in R for the analysis is available in this paper’s supplemental information. The remaining data reported in this paper will be shared by the lead contact upon request.

### METHOD DETAILS

#### Co-immunoprecipitation assay

Brains from C57BL/6 mice (Janvier Labs) or aSN-KO (Sncatm1Rosl (C57BL/6), The Jackson Laboratory) were homogenized in 7x w/v homogenization buffer (320 mM sucrose, 4 mM HEPES–NaOH, 2 mM EDTA, and Complete protease inhibitor mix (Roche), pH 7.4) using a loose-fitting glass-Teflon homogenizer (10 up-and-down strokes, 700 rpm). Debris was removed from the homogenate by centrifugation for 10 min at 1000 x g in a Sorvall RC 5C plus centrifuge. The resulting supernatant was centrifuged for 1 h at 100 000 x g. The supernatant was removed, and the pellet was resuspended in the original volume (7x) of RIPA (50 mM Tris pH 7.4, 159 mM NaCl, 1% Triton X-100, 2 mM EDTA, 0.5% sodium deoxycholate, 0.1% SDS) for 3 h, whereafter samples were spun at 20 000 x g, for 30 min at 4°C. Protein concentration was measured using the bicinchoninic acid assay. aSN oligomer or monomer (2 μg/ml) was mixed with 0.5 mg/ml aSN-KO mouse brain homogenate diluted in PBS, 0.5% Triton X-100 and incubated overnight at 4°C. aSN binding (ASY-1) or control (non-immune IgG) was performed as previously described (Betzer *et al*., 2015). The samples were incubated for 2 h with rotation. The Sepharose beads were isolated and washed twice with PBS, 0.5% Triton X-100, and Co-IP proteins were eluted by incubation in a non-reducing SDS loading buffer at room temperature. Proteins were resolved on 10–16% gradient SDS-PAGE under reducing conditions followed by immunoblotting for PMCA (primary antibody: 5F10 anti-pan-PMCA, Abcam, ab2825, secondary antibody: anti-mouse-HRP, Dako) and aSN (primary antibody: anti-Syn-1, BD Transduction Laboratory, secondary antibody: anti-mouse-HRP, Dako). The interaction between endogenous aSN and the endogenous PMCA was studied in the extracts from C57BL/6 mice as described above.

#### Primary hippocampal neuronal cultures and cytosolic Ca^2+^ measurements

Primary hippocampal neurons were cultured from newborn (P0) aSN-KO mice (Sncatm1Rosl (C57BL/6), The Jackson Laboratory). Hippocampi were dissected in ice-cold Hank’s balanced salt solution, dissociated in 20 U/ml papain in Hibernate A medium (Gibco) supplemented with 1xB27 and 0.3 g/L l-glutamine for 20 min at 37°C, washed twice, and triturated in plating medium [MEM (Gibco) supplemented with 5 g/l glucose, 0.2 g/l NaHCO_3_, 0.1 g/l transferrin, 0.25 g/l insulin, 0.3 g/L l-glutamine, and 10% fetal bovine calf serum (heat-inactivated)]. Hippocampal neurons were seeded on Matrigel® matrix (Corning ®)-coated coverslips. After 24 h, the medium was changed to growth medium [MEM supplemented with 5 g/l glucose, 0.2 g/l NaHCO_3_, 0.1 g/l Transferrin, 0.075 g/L l-glutamine, 1×B-27 supplement, 2 µM cytosine arabinoside and 5% fetal bovine calf serum (heat-inactivated)].

At culture day 6 (DIV 6), the neurons were transfected by lipofectamine 3000 with Bicistronic vectors coding for mCherry and aSN or mCherry under synapsin promotor according to manufacturer’s instructions. At culture day 8 cytosolic Ca^2+^ levels in primary neurons were determined using the Ca^2+^-sensitive fluorescent indicator Fura-2-AM (Molecular Probes/Invitrogen). Cells were loaded with Fura-2 in sterile-filtered HEPES-buffered saline (HBS: 20 mM HEPES, 150 mM NaCl, 5 mM KCl, 1 mM CaCl_2_, 1 mM MgCl_2_, 10 mM glucose, pH 7.4) containing 2.5 μM Fura-2 AM, 0.04% pluronic acid, F127 for 30 min at 37°C, 5% CO_2_. The Fura-2-containing medium was replaced with fresh HBS without Fura-2 and incubated additionally for 30 min. After dye loading and prior microscopic analysis, the culture was moved into a recording buffer with thapsigargin for 5 min., and an area containing 1-3 transfected neurons was found. The fluorescence was measured on an Olympus Scan^R high-content microscope using excitation wavelengths at 340 and 380 nm and emission at 510 nm. The cytosolic Ca^2+^ levels in single cells were measured by placing a region of interest (ROI) outside the nucleus. The recording was started and at the recording time of 2 min KCl was added to depolarize the neurons. The calcium response was followed over time until 12 min. The response upon K^+^ induced depolarization was quantified as the Area Under Curve (AUC) from each measured neuron from the 2-12 min. measurement interval.

#### Proximity ligation assay (PLA)

Primary hippocampal neurons were cultured from newborn (P0) C57BL/6 mice (Janvier Labs) as described for aSN-KO. At culture day 14 (DIV14) the neurons were fixed in 4% paraformaldehyde for 30 min at room temperature (RT) followed by a wash in PBS and 10 min permeabilization in 0.1% Triton X□100, 50 mM glycine, 3 mM CaCl_2_, 2 mM MgCl_2_, pH 7.4. Unspecific binding was blocked by 3% bovine serum albumin in PBS for 1 h followed by incubation with the primary antibody for 1.5 h [anti□AS (1 μg/ml, ASY□1(Kragh *et al*, 2009; Lindersson *et al*, 2004)), and anti□PMCA 5F10 (1 μg/ml, Abcam, ab2825). The Duolink procedure was conducted according to the manufacturer’s instructions with the Duolink® *In Situ* Red Starter Kit Mouse/Rabbit (Duolink®, Sigma□Aldrich), with secondary antibodies contained in the kit. After PLA staining, the neurons were labeled with synaptophysin (primary antibody - guinea pig anti-synaptophysin 1, (Synaptic Systems #101004), secondary: goat anti-guinea pig, Alexa Fluor® 488 nm (Abcam, ab150185) to visualize the synapses where synuclein normally is located and DAPI for nuclei. Images were obtained using a Zeiss Observer Z1 inverted microscope equipped with ApoTome.2.

#### Expression and purification of α-synuclein

Recombinant human aSN wild type and variants were produced in *E. coli*, and purified as previously described (Huang *et al*, 2005). Monomeric and oligomeric forms of aSN were produced and isolated as previously described (Betzer *et al*., 2015). Concentration of all aSN preparations was confirmed by BCA assay. The proteins were stored in -80□ in a buffer containing 20 mM Tris/HCl, pH 7.5, 150 mM KCl. Recombinant ^15^N-aSN for NMR experiments and aSN(Δ1-14) was prepared as previously described (Cholak *et al*., 2020)(PMID: 32277854) and aSN(1-61) as in (Skaanning *et al*., 2020).

#### PMCA expression constructs and site-directed mutagenesis

Codon-optimized genes encoding for human PMCA1d and PMCA2w/a (Genscript) were cloned into pEMBL-yex4 expression plasmids by homologous recombination in *S. cerevisiae* as described (Drew *et al*, 2008).

PMCA1d and PMCA2w/a constructs were subjected to site-directed mutagenesis using QuikChange site-directed mutagenesis kit according to the manufacturer’s protocol (Agilent). To obtain an inactive variant D475N (the autophosphorylation site) mutagenetic primers were designed using PrimerX online tool (https://www.bioinformatics.org/primerx). To obtain C-terminally truncated PMCA1, an internal TEV cleavage site was introduced to the original construct. Primers for the insertion of codon-optimized TEV site sequence and the large deletions in PMCA2 were designed according to described methods (Liu & Naismith, 2008). The integrity of all constructs was verified by DNA sequencing (Eurofins).

#### Expression of recombinant PMCAs in *S. cerevisiae*

All circular vectors were introduced into yeast cells using PEG/LiAc/ssDNA-mediated transformation (Gietz & Schiestl, 2007). All recombinant PMCAs were expressed in *S. cerevisiae* strain K616, depleted of the native calcium ATPases (*MATα pmr1::HIS3 pmc1::TRP1 cnb::LEU2 ura3-1*) (Cunningham & Fink, 1994). All culture media were supplemented with 10 mM CaCl_2_, as it is a necessary survival condition for the strain to maintain internal stores. For expression culture, 100 ml of uracil-deficient 2% glucose SD medium was inoculated with a colony of transformed cells and allowed to grow for 24 h in 30°C in a 120 rpm shaking incubator. The preculture was used for inoculation of 9 L of -Ura SD medium with 1% glucose and 40 mg/L adenine hemisulfate, starting from OD_450_ of 0.05. After 22 hours, when all glucose in the medium was used and the culture reached OD_450_ of ∼5, the expression was induced by 2% galactose, and the culture was supplemented with YP medium and 40 mg/L adenine hemisulfate. Cells were harvested after 18-20 h by centrifugation, washed with water and TEKS buffer (50 mM Tris/HCl pH 7.6, 2 mM EDTA, 100 mM KCl, 0.6 M D-sorbitol), and resuspended in TESin buffer (50 mM Tris/HCl, 2 mM EDTA, 0.6 M D-sorbitol with the addition of PMSF and protease inhibitor. The typical yield was ∼15 g of cells per liter of culture.

#### Purification of PMCAs

The cells were disrupted in a pulverisette grinder with an equal volume of 0.5 mm glass beads (4°C, 450 rpm, 4 cycles of 3 min millings, and 1 min pause). The material was centrifuged for 20 min at 2000 x g to remove beads and uncracked cells. The supernatant (S1) was centrifuged for 30 min at 20 000 x g for further removal of cell debris. The supernatant (S2) was subjected to ultracentrifugation for 3 h at 150 000 x g. Pelleted membranes were suspended in solubilization buffer (20% glycerol, 50 mM Tris/HCl pH 7.2, 150 mM KCl, BME, PMSF, protease inhibitors) and homogenized in a Dounce glass homogenizer. Membranes were flash-frozen in liquid nitrogen and stored at -80°C until further processing.

Membranes were diluted with solubilization buffer to 5 mg/ml total protein concentration (measured with Bradford assay) and solubilized in 1% DDM and 0.2% cholesteryl hemisuccinate for 1 h by stirring in 4°C, followed by ultracentrifugation at 150000 x g for 1 h to pellet unsolubilized material. The supernatant was supplemented with 4 mM CaCl_2_ and stirred for 2 h in 4°C with CaM-Sepharose beads (GE Healthcare), pre-equilibrated with binding buffer (50 mM Tris/HCl pH 7.2, 150 mM KCl, 0.017% w/v DDM, 10% glycerol, 2 mM CaCl_2_). The beads were packed onto a gravity flow column and washed with 10-15 column volumes of binding buffer. The protein was eluted a buffer containing 50 mM Tris/HCl pH 7.2, 150 mM KCl, 0.017% w/v DDM, 10% glycerol, 2 mM EGTA. EGTA stock solution (200 mM) was beforehand pH-adjusted with KOH to ∼7.5. The fractions containing the protein of interest were pooled and concentrated to ∼3 mg/mL using a 100kDa MWCO Vivaspin protein concentrator (GE Healthcare). The concentrated protein was subjected to size exclusion chromatography on Superose 6 increase 10/300 column (GE Healthcare).

To produce C-terminally truncated PMCA1, the PMCA1d construct with a tobacco etch virus (TEV) protease cleavage site inserted next to H1074 just before the autoinhibitory domain was used. The protein was cleaved with TEV protease while bound to the CaM sepharose beads. After overnight incubation with the protease at 4□, the C-terminally truncated PMCA was collected in the flow-through fraction and run over a Ni^2+^ affinity column to bind the His-tagged protease. The flow-through was concentrated to ∼3mg/mL (100kDa MWCO) and subjected to size exclusion chromatography on Superdex 200 increase 10/300 column (GE healthcare) in size exclusion buffer containing 50 mM Tris/HCl pH 7.2, 150 mM KCl, 0.017% w/v DDM, 10% glycerol. Peak fractions of PMCAs were pooled, concentrated to 1-1,5 mg/ml if necessary, using a 100kDa MWCO Vivaspin protein concentrator (Sartorius), aliquoted, and flash-frozen in liquid N_2_ and stored at -80°C for further procedures. The purity of the proteins was analyzed by SDS-PAGE (Supplementary Fig. 1).

#### Expression and purification of CaM

Mammalian CaM in the pET15b vector was transformed into E. coli strain C41 and grown in LB media at 37°C. Induction was performed with IPTG at an OD_600_ of 0.6. After 16 hours of growth at 21°C, cells were harvested by centrifugation at 3000 x g, resuspended in lysis buffer (20 mM Tris-HCl pH 7.5, 1 mM EDTA, cOmplete protease inhibitor), and lysed using sonication. The lysate was centrifuged for 1 h at 20 000 x g. The supernatant was mixed 1:1 with wash buffer (20 mM Tris-HCl pH 7.5, 10 mM CaCl_2_) and loaded onto a pre-equilibrated (wash buffer) gravity flow phenyl sepharose column. The column was washed with 5 CV wash buffer, 5 CV high salt wash buffer (20 mM Tris-HCl pH 7.5, 500 mM NaCl, 10 mM CaCl_2_), and 5 CV wash buffer. CaM was eluted in elution buffer (50 mM Tris-HCl pH 7.5, 150 mM NaCl, 10 mM EGTA). Before use in activity assays, CaM buffer was exchanged to 20 mM Tris-HCl pH 7.5, 150 mM KCl using PD MiniTrap G25 desalting column (GE Healthcare), aliquoted and stored in -80□.

#### Expression and purification of ACA8

ACA8 expression and membrane isolation was performed similarly to the PMCA isoforms. ACA8 was solubilized in a DDM solubilisation buffer (50mM Tris-HCl pH 7.6, 100mM KCl, 20% glycerol, 3mM MgCl_2_, 5mM β-mercaptoethanol (βME), 7.5 mg/mL DDM, 1mM PMSF and 1µg/mL of chymostatin, pepstatin A and leupeptin). Purification was achieved through Ni^2+^ affinity followed by size exclusion chromatography. ACA8 was bound to a Ni^2+^-affinity column in low imidazole buffer (50 mM Tris-HCl pH 7.5, 10 mM imidazole, 200 mM KCl, 10 % glycerol, 3 mM MgCl_2_, 5 mM βME, 0.2 mM LMNG). On the column, the ACA8 His-tag was cleaved off with thrombin and ACA8 was eluted in low imidazole buffer. ACA8 was subjected to size exclusion chromatography on a Superose 6 increase 10/300 column in a SEC buffer (50 mM Tris-HCl pH 7.5, 200 mM KCl, 3 mM MgCl_2_, 10% glycerol, 0.02 mM LMNG, 5 mM βME). The protein was stored in -80□ in SEC buffer for further experiments.

#### Expression and purification of PMCA2w-ATM3-loop

*E. coli* cells (BL21 DE3) were transformed with a modified pET24b plasmid containing N-terminal His_6_-SUMO tagged PMCA2w_A-TM3_ loop (L289-K383) using heat shock transformation. Pre-cultures (10 mL LB medium, 50 μg/ml kanamycin) were grown overnight and used to inoculate 1 L of ZYM-5052 media (PMID: 15915565), supplemented with 50 μg/ml kanamycin. Cells were grown for 3 h at 37 □ with shaking, then transferred to a shaking incubator at 16 □ and grown for a further 24 h. The cells were pelleted at 4,000 × *g*, and kept at -20 □ for at least 24 h prior to lysis in column binding buffer (50 mM Tris pH8, 150 mM NaCl, 10 mM Imidazole), supplemented with cOmplete protease inhibitor (Sigma), at 20 kpsi using a French pressure cell disruptor (Constant Systems, Daventry, UK). Lysate was cleared by centrifugation at 20,000 × *g*, then passed through a 0.45 μm filter before being incubated with equilibrated Ni-Sepharose fast flow resin (GE Healthcare) for 30 min. Lysate was passed through the column via gravity flow, followed by wash buffer (5 × CV) (50 mM Tris pH8, 1 M NaCl, 10 mM Imidazole) 5 × CV binding buffer, and eluted with elution buffer (10 mL; 50 mM Tris pH 8, 150 mM NaCl, 250 mM Imidazole). Purified His_6_-SUMO-PMCA2w was made up to 50 mL (with 50 mM Tris pH 8, 150 mM NaCl) and cleaved with ULP1 (0.1 mg) ON at 4 □. The cleaved protein was purified by collecting the flow-through from a second Ni-Sepharose column and concentrated using Amicon spin filters (Millipore). Protein was stored at -20 □.

#### NMR experiments

All NMR spectra were recorded at 5°C on a Bruker 800 MHz spectrometer equipped with a cryogenic probe and Z-field gradient using Bruker Topspin v4.0.7, recording ^15^N,^1^H--HSQC spectra to address interaction and using the backbone assignments obtained in (Cholak *et al*., 2020). ^15^N-HSQC spectra were obtained in HEPES buffer (20 mM HEPES pH 7.4, 150 mM NaCl) of 50 *μ*M ^15^N-aSN alone and in the presence of PMCA2w_ATM3_-loop (molar ratio 1:5) and with further addition of 25 mM CaCl_2_ (molar ratios 1:4:500). Spectra of 9.5 *μ*M ^15^N-aSN alone and with full-length PMCA2w/a (molar ratio 1:1) were obtained in Tris buffer (50 mM Tris pH 7.5, 150 mM KCl, 1.5 % (w/v) DDM). Combined amide (N, H^N^) chemical shift perturbation (CSPs) were measured by CCP-analysis version 2.5. The spectral analyses were done in CcpNMR analysis (Vranken *et al*, 2005).

#### Far-UV CD spectropolarimetry

Far UV CD spectra were recorded at 5°C on a JASCO J-815 spectropolarimeter on 12.8 *μ*M PMCA2w_A-TM3_ loop dissolved in 10 mM Na_2_HPO_4_, 200 mM NaF pH 7.4. The spectrum was recorded from 260-190 nm, averaging 10 scans (D.I.T 2 s; data pitch 0.2 nm; band width 1 nm; scan rate 50 mm/min; path length 1 mm). A spectrum of the buffer was recorded using identical settings and subtracted.

#### SERCA and Na, K-ATPase preparations

SERCA1a from rabbit skeletal muscle was kindly donated by Thomas Lykke-Møller Sørensen, Anne Lillevang and Claus Olesen. The protein preparation was performed according to previously established procedures (Andersen *et al*, 1985; Sorensen *et al*, 2004). Na, K-ATPase was kindly donated by Michael Habeck and Natalya Fedosova. The protein was expressed and purified as described earlier (Habeck *et al*, 2017)

#### ATPase activity assay

The ATPase activity was tested by monitoring the release of free inorganic phosphate from ATP in a molybdenum blue assay (Baginski *et al*, 1967).

Purified PMCA requires relipidation to regain activity, therefore the samples were preincubated with phospholipids before activity assay. Lipid stock solutions were prepared from powdered lipids as follows: brain lipid extract (from bovine brain Type I, Folch Fraction I, Sigma-Aldrich), brain PC (Avanti), or soy PC (Avanti) were dissolved to a concentration of 20 mg/ml in a buffer containing 50 mM Tris pH 7.5, 150 mM KCl and 1.5 % (w/v) DDM. After cycles of heating (45-50□), sonicating, and vortexing, a translucent gel was obtained. Stocks were aliquoted and stored in -20°C. Shortly before the assay lipid solution was added to PMCA sample in PMCA:lipid w/w ratio 1:5 and mixed by gentle pipetting. After 15 min, the relipidated protein was added to a final concentration of 2.5 μg/ml to a reaction buffer containing 50 mM KNO_3_, 5 mM NaN_3_, 0.25 mM NaMob, 40 mM BisTris/HEPES pH 7.2, 3 mM MgSO_4_, 1 mM EGTA and CaCl_2_ in concentration needed for obtaining the desired concentration of free calcium ions. The free calcium concentration was calculated using Maxchelator Ca/Mg/ATP/EGTA Calculator v1.0 using constants from NIST database (https://somapp.ucdmc.ucdavis.edu/pharmacology/bers/maxchelator/CaMgATPEGTA-NIST.htm) aSN and/or CaM were added to the reaction mixture in desired concentrations and corresponding protein storage buffer was added to control samples in equal volume. The reactions were started by adding 3 mM ATP, carried out at 37°C, and stopped after 7 or 11 min by adding a stop solution. Stop solution was prepared shortly before every assay by mixing ice-cold solutions A (17 μM ascorbic acid, 0.1% SDS in 0.5 M HCl) and B (0.7 μM ammonium molybdate VI tetrahydrate). The absorbance was measured with Victor 3 96-well plate reader (Perkin-Elmer) at 860 nm.

#### Transcriptomics data analysis

Data on gene expression for PMCA1-4 and aSN were used. Data on tissue expression and exon incorporation from VastDB for the genes ATP2B1, ATP2B2, ATP2B3, ATP2B4, and SNCA were imported in R. For the PMCA exons of relevance, the “percent spliced-in” (PSI) was correlated to aSN tissue expression. For tissues in specified aSN expression intervals (ranging from 0-190 interval size of 10) the average incorporation of the exons was determined, and the mean PSI was plotted as a function of the aSN expression interval. The code written in R for the analysis is included in the supplemental information.

#### Protein disorder prediction

Protein disorder prediction was performed using NetSurfP 2.0 software developed by the Technical University of Denmark and available at https://services.healthtech.dtu.dk/service.php?NetSurfP-2.0.

#### Statistical analysis

Statistics were performed using GraphPad Prism and P values <0.05 were considered significant. The statistical method used is described in the figure legends.

## SUPPLEMENTAL INFORMATION TITLES AND LEGENDS

**Figure S1.**
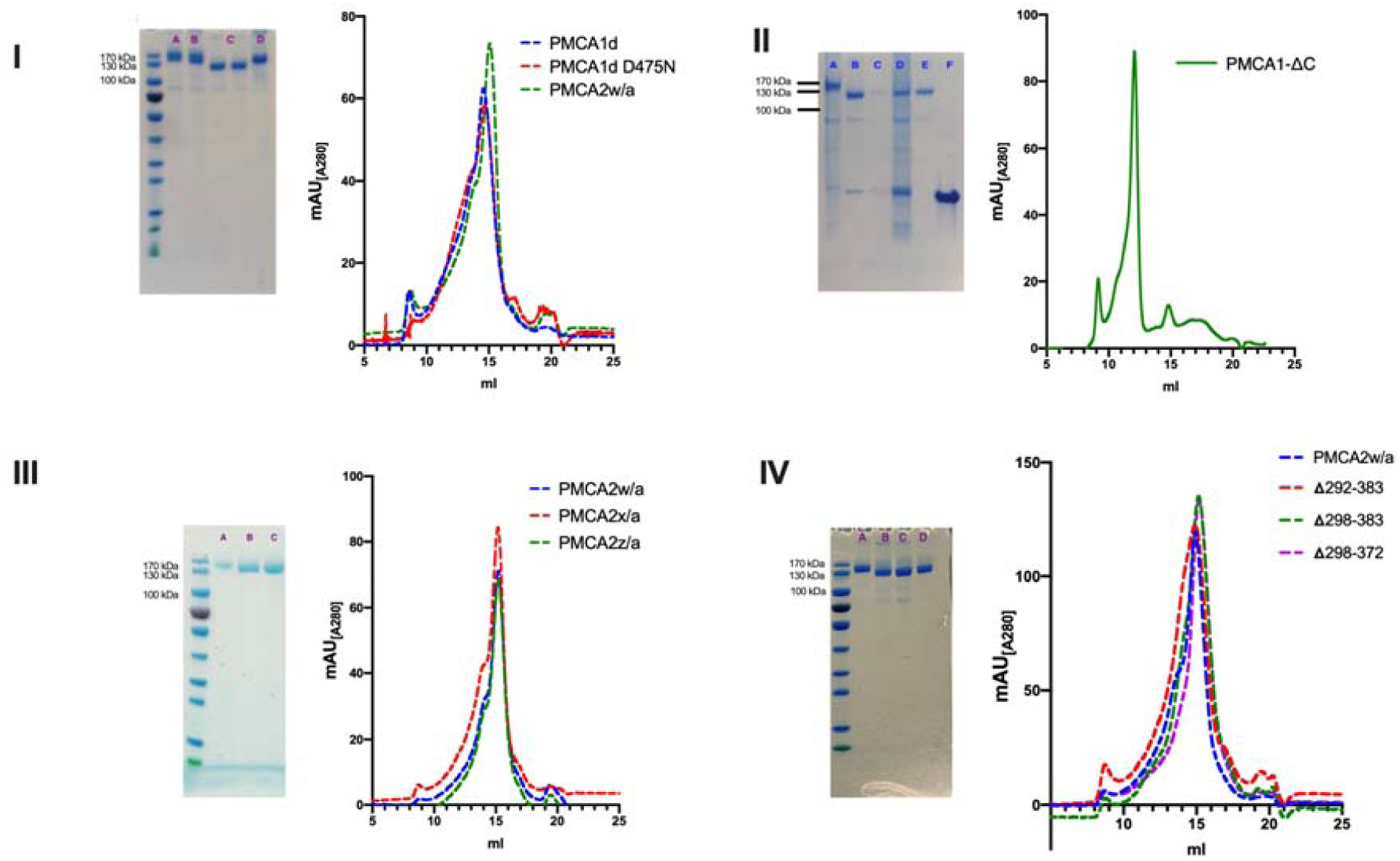
Purification of recombinant PMCAs. **I** - *Left* - SDS PAGE of purified PMCAs. (A) PMCA1d, PMCA1d D475N (inactive variant), PMCA1-ΔC, (D) PMCA2w/a. *Right* **–** Corresponding profiles of the size exclusion chromatography performed on the Superose 6 increase column. **II** – *Left* – SDS-PAGE analysis of the purification of the PMCA1-ΔC. Full PMCA1d with internal TEV cleavage site at position 1074 was bound to CaM sepharose and washed (lane A – a sample of washed beads). After overnight incubation with His-tagged TEV flow-through 1 (FT1, lane B) was collected and beads were washed with buffer in a gravity-flow column (lane C - wash). FT1 was put through a Ni-Sepharose in a gravity-flow column to bind TEV and uncleaved PMCA1 (D – sample of washed sepharose beads). The flow-through from the nickel column was collected, concentrated to 500 µL, and subjected to size exclusion chromatography on Superdex 200 column. Lane E – peak fraction from the size-exclusion chromatography. Lane F – TEV protease. *Right* – Profile of the size exclusion chromatography of PMCA1-ΔC performed on the Superdex 200 increase column. **III** – *Left* - SDS PAGE of purified PMCA2 splice variants (A-C) w/a, x/a, z/a. *Right* **–** Corresponding profiles of the size exclusion chromatography performed on the Superose 6 increase column. **IV** - *Left* - SDS PAGE of purified PMCA2w/a (A) and its deletion variants Δ292-383 (B), Δ298-383 (C), and Δ298-372 (D). *Right* **–** Corresponding profiles of the size exclusion chromatography performed on the Superose 6 increase column.

**Figure S2.**
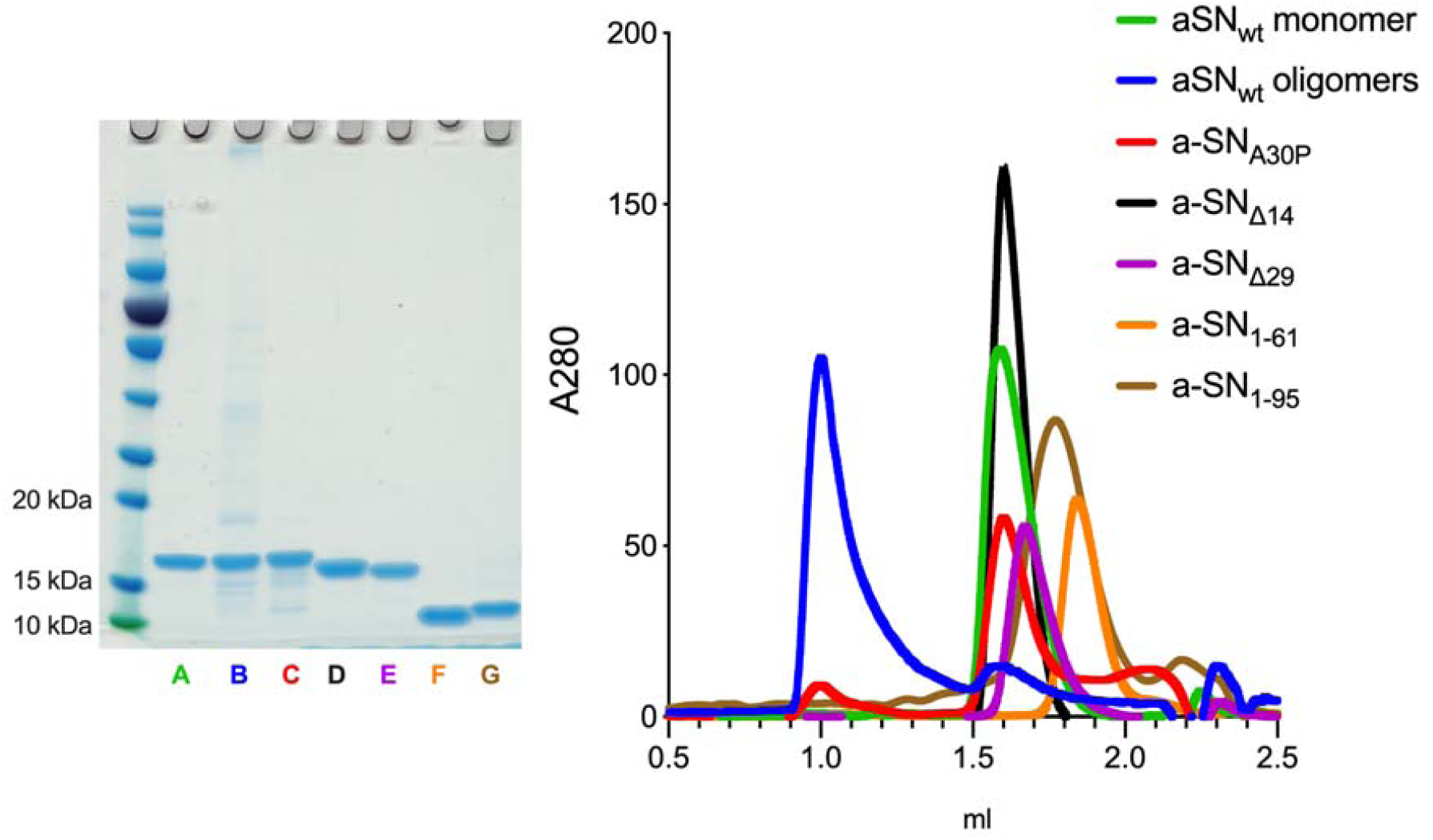
Purity of alpha-synuclein preparations. *On the left* – SDS-PAGE analysis A – monomer; B – oligomers; C – A30P; D – Δ14; E – Δ29; F – (1-61); G – (1-95); *On the right* – size exclusion chromatography analysis performed on Superdex 200 increase 3.2/300 column. Monomeric forms were eluted at volume > 1.5 ml

**Figure S3.**
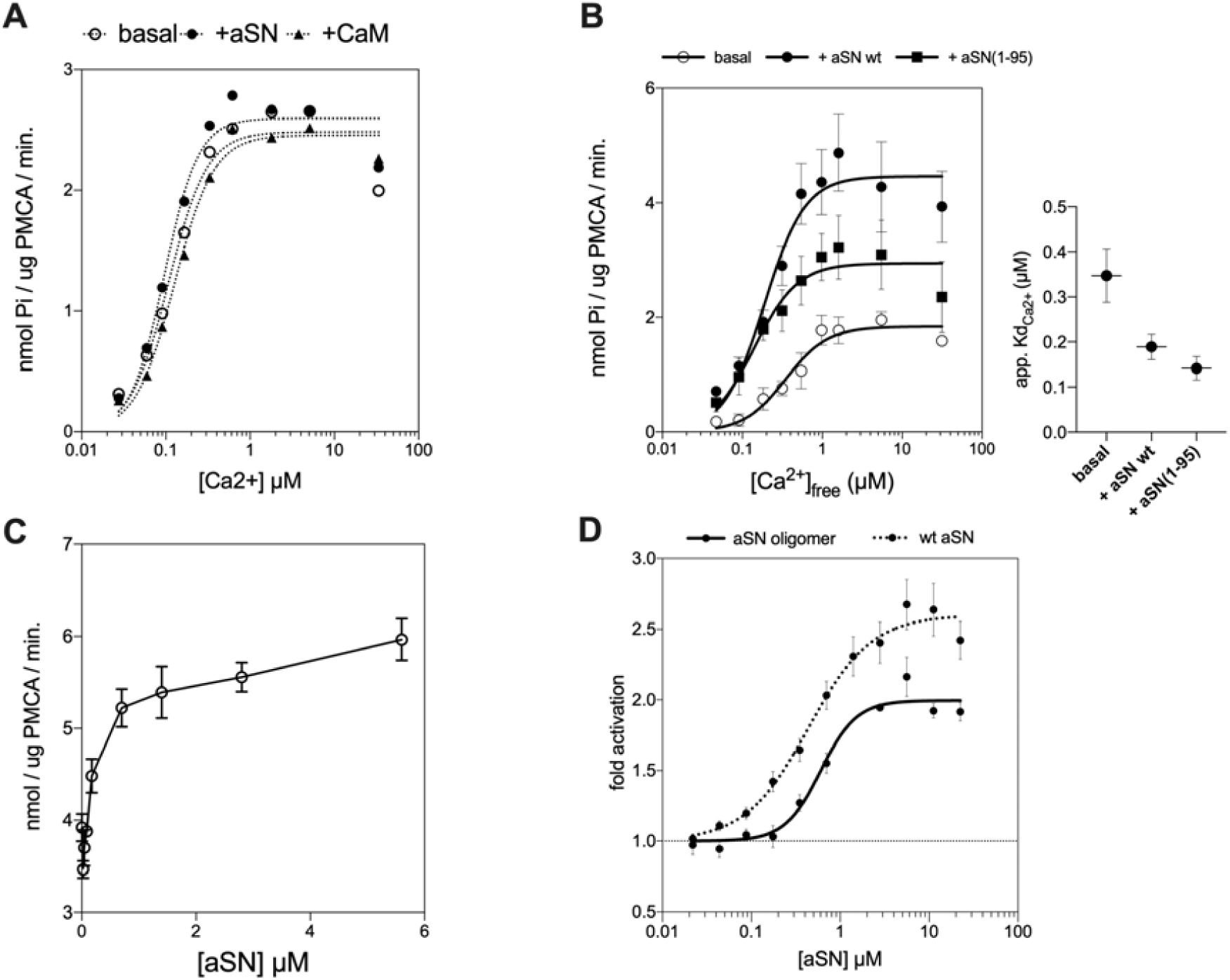
**A - The activity of PMCA1-**Δ**C in the presence of brain PC lipids**. Ca^2+^ titrations were performed in absence of an activating partner, in the presence of 2.8 μM alpha-synuclein or 1.2 μM CaM. **B - The effect of full-length and C-terminally truncated aSN**_**(1-95)**_ **on the calcium-dependent activity of PMCA1d**. *On the left* – the calcium titration experiment was performed simultaneously in absence of activating partner (*empty circles*), in presence of the full-length monomeric a-SN (*filled circles*) and C-terminally truncated aSN_(1-95)_ (*filled squares*). The pump was relipidated in brain lipid extract (BE). *On the right* – apparent Kd_Ca2+_ values calculated from the fitted Hill plots. The lines are the best fit given by the Hill equation with the apparent Kd values (μM) for Ca^2+^ as follows: PMCA1d basal (without aSN) – 0.347 ± 0.059, with aSNwt - 0.189 ± 0.028, with aSN_(1-95)_ – 0.142 ± 0.026 **C -** The activity of C-terminally truncated PMCA4x (lacking the autoinhibitory domain) was measured as a function of monomeric a-SN concentration in presence of 1.8 μM free Ca^2+^. PMCA was relipidated with brain extract (BE). **D - The effect of oligomeric aSN on the PMCA activity**. PMCA1d fold activation by titrated full-length monomeric and oligomeric aSN. Monomeric aSN data, presented here for comparison, is the same as in Figure 4. The assay was performed in presence of 1.8 μM free Ca^2+^ and brain lipid extract was used for the PMCA relipidation. Data are a result of at least three independent activity assays; mean ± SEM.

**Figure S4.**
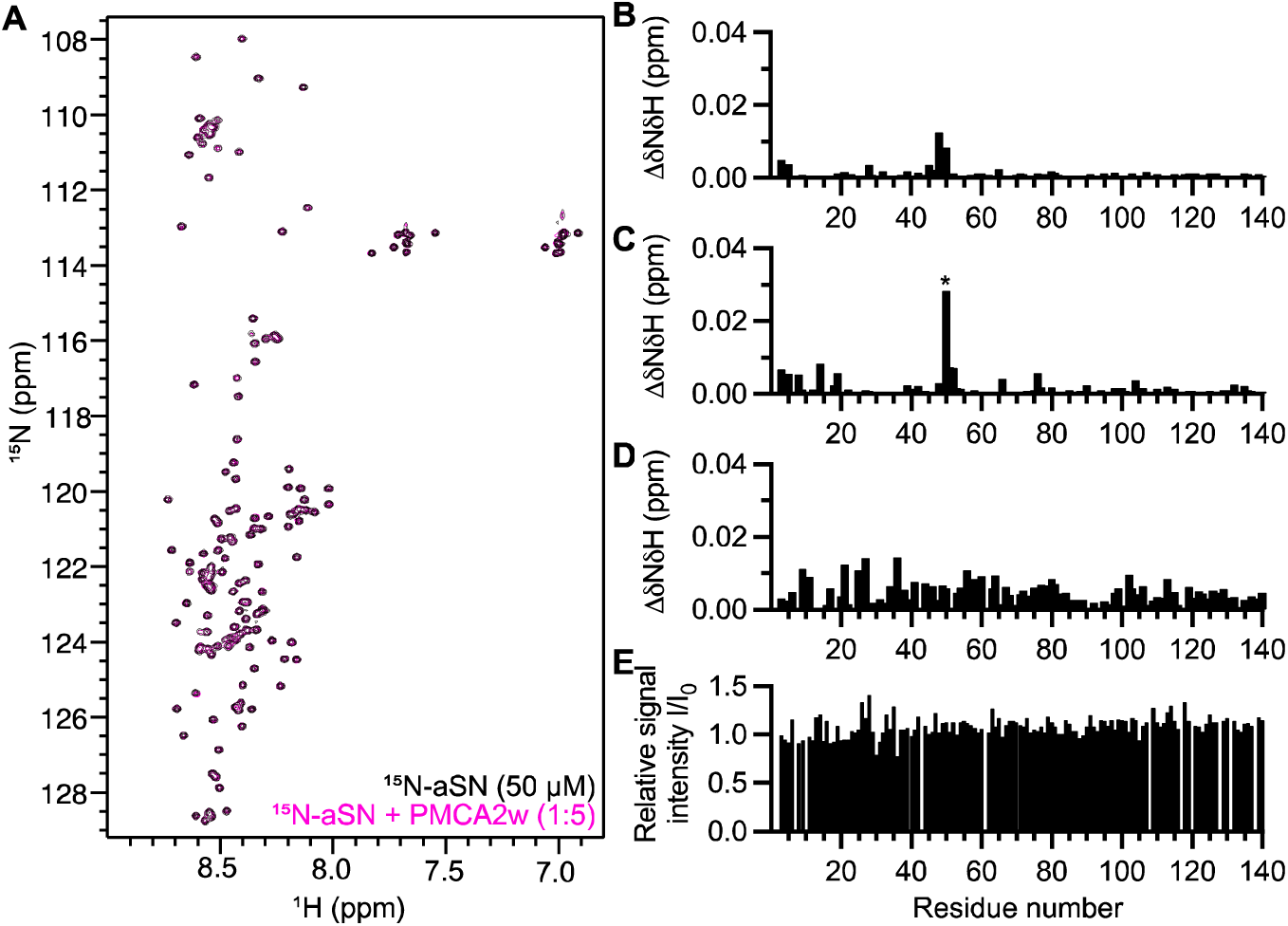
NMR interaction studies. **A-** ^15^N-HSQC spectra comparing ^15^N-aSN alone (black) with that of ^15^N-aSN added PMCA2w_A-TM3_ loop (pink). **B-** Quantified amide chemical shift perturbations (CSPs) from **A**. **C-** Amide CSPs caused by addition of PMCA2w_A-TM3_ loop to ^15^N-aSN in the presence of calcium (^15^N-aSN:Ca^2+^ 1:500; ^15^N-aSN:Ca^2+^:PMCA2w 1:500:4). * indicates pH sensitivity observed for histidine. **D-** CSPs caused by full-length PMCA2w/a at a 1:1 molar ratio with ^15^N-aSN. **E-** Peak intensity changes caused by addition of full-length PMCA2w/a at a 1:1 molar ratio with a^15^N-aSN.

**Figure S5.**
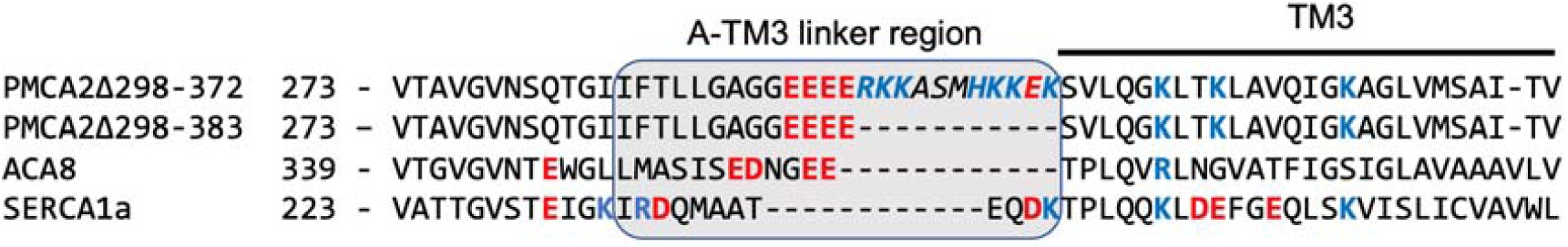
Sequence alignment of the A-TM3 loop region and the TM3 domain of PMCA2w/aΔ298-372, PMCA2w/aΔ298-383, ACA8 and SERCA1a. Positively charged residues marked blue, negatively – red. Italics show the acidic lipid binding site of PMCA2. Alignment of full sequences was performed with uniport.org, sequences from which presented fragments were taken have the following entry numbers in Uniprot: Q01814-2 (PMCA2w/a, modified for the alignment, to show both PMCA deletion variants), Q9LF79 (ACA8), and P04191 (SERCA1a).

